# Mediodorsal thalamus regulates sensory and mapping uncertainties in flexible decision making

**DOI:** 10.1101/2022.12.11.519975

**Authors:** Xiaohan Zhang, Michael M. Halassa, Zhe Sage Chen

## Abstract

The mediodorsal (MD) thalamus is a critical partner for the prefrontal cortex (PFC) in cognitive flexibility. Accumulating evidence has shown that the MD regulates task uncertainty in decision making. However, the mechanism of this cognitive process remains unclear. Here we used a reverse-engineering approach and trained biologically-constrained computational models to delineate these mechanisms. We found that the inclusion of an MD-like feedforward module increased robustness to sensory noise, enhanced working memory and enabled rapid context switching in the recurrent PFC network performing two versions of context-dependent decision-making tasks with sensory and mapping uncertainties. Incorporating genetically identified thalamocortical pathways and interneuron cell types replicated neurophysiological findings of neuronal tuning and uncovered attractor-like population dynamics. Our model revealed key computational mechanisms of context-invariant MD in regulating cueing uncertainty and context switching. It also made experimentally testable predictions linking cognitive deficits with disrupted thalamocortical connectivity, prefrontal excitation-inhibition imbalance and dysfunctional inhibitory cell types.

## INTRODUCTION

Behavioral flexibility and cognitive control are fundamental in decision making;^1^ successful execution of complex decision-making tasks requires identification and processing of multiple sources of uncertainty^2^. Task uncertainty may appear in the form of corrupted or incongruent sensory cues (“sensory uncertainty”)^3,4^, their mapping onto internal or behavioral variables (“mapping uncertainty”)^2,5,6^, or their likelihood of resulting in reward (“outcome uncertainty”)^7^. Behavioral flexibility to map the same rule under different contexts is critical in decision making. Experiments across multiple species have shown that the MD thalamus is an important partner for the PFC in resolving uncertainty in decision making^8-15^. Human neuroimaging studies have shown that MD activity tracks sensory uncertainty in a multi-attribute attention task^14^ and a categorization task^16^. This process generalizes to non-human animals; in mice performing a decision-making task, the MD tracks sensory uncertainty and enables a rapid switching of cue-to-rule transformation^6,17^. Optical manipulations support this notion and delineate their causal roles in cognitive control^5,18^. However, the computational mechanism by which the MD enhances prefrontal activity in context-dependent decision making under sensory and mapping uncertainties remain unclear. In addition, how the newly discovered cellular diversity in MD thalamus contributes to such computations is unexplored. Biologically-inspired computational modeling that incorporates neuronal subtype and circuit pathway knowledge^19-21^ may provide an approach to probe these questions and yield mechanistic insight^22-29^.

In light of the tenet “structure determines functions”, an important circuit computation question arises: how does a feedforward MD-like structure facilitate neural computation in a recurrent PFC network to regulate different task uncertainties? More specifically, how cortical interneuron cell types, intracortical and thalamocortical connectivity may influence task performance and generate emergent task-specific neural representations? To investigate these questions, here we used a reverse-engineering approach and trained biologically-constrained PFC-MD models to perform two versions of context-dependent decision-making tasks, one with sensory uncertainty and the other with both sensory and mapping uncertainties, from which we delineated the roles of interneuron subtype and thalamocortical-projection specificity in prefrontal computation. Our PFC-MD model replicated key behavioral data and neuronal tunings in task-performing mice^5,6^, including prefrontal context-invariant rule-tuning and thalamic context tuning, and further made new experimentally testable predictions. The MD subpopulations and cell-type specific MD-to-PFC projections had distinct functional roles in regulating sensory uncertainty driven by cue conflict and sparsity, maintaining working memory, and mediating prefrontal computation in a task-phase specific manner. To accommodate mapping uncertainty, synaptic plasticity of thalamocortical connections enabled rapid context switching to perform rule-invariant prefrontal computation. Our analysis suggests that the feedforward MD regulates recurrent prefrontal computation to improve information integration by increasing intrinsic time constant and controls cue-to-rule remapping by thalamocortical plasticity. Furthermore, the PFC-MD models enabled us to parse cognitive deficits of thalamocortical circuits in flexible decision making with task uncertainty, which are induced by prefrontal excitation-inhibition (E/I) imbalance, dysfunctional inhibitory cell types, and disrupted thalamocortical and corticothalamic connectivity.

## RESULTS

### Training biologically constrained PFC-MD models to perform decision-making tasks with parameterized sensory uncertainty

We adopted a task-optimized training strategy of computational models to delineate the computational mechanism of context-dependent decision making. We first modeled the PFC with a rate-based excitatory-inhibitory (E/I) recurrent neural network (E/I-RNN) (Methods), where cortical GABAergic inhibitory neurons consisted of parvalbumin (PV), vasoactive intestinal peptide (VIP)-expressing and somatostatin (SOM) interneuron cell types (**Fig. 1a**). We assumed structured inhibitory-to-excitatory and inhibitory-to-inhibitory connectivity and disallowed weak connections that are negligible. Specifically, the VIP interneurons mainly disinhibit other classes of interneurons^30,31^. We imposed several key biological constraints on network connectivity. First, we modeled the MD as a feedforward structure devoid of lateral connectivity, where all MD neurons received excitatory inputs from the PFC and projected back to excitatory neurons. Second, guided by recent knowledge of genetically identified thalamocortical projections that differentially target distinct PFC interneuron types^5^, we imposed additional connectivity constraints onto our model. One MD subpopulation, which is termed as MD_1_, projects to prefrontal PV interneurons, while the other subpopulation, MD_2_, projects to cortical VIP interneurons. To translate them to biological terms, MD_1_⟶PV corresponds to the kainite receptor (GRIK4)-expressing (i.e., MD_GRIK4_) pathway with preferentially targeted PV+ neurons; MD_DRD2_⟶VIP projection corresponds to the dopamine receptor (D2)-expressing (i.e., MD_DRD2_) pathway with preferentially targeted VIP^+^ neurons. We trained both the PFC-MD model, and a “PFC-alone” model as a control (using identical stimulus inputs and cortical E/I configuration) to perform a cueing context-dependent decision-making task (**Fig. 1b** and Methods).

**Figure 1.**
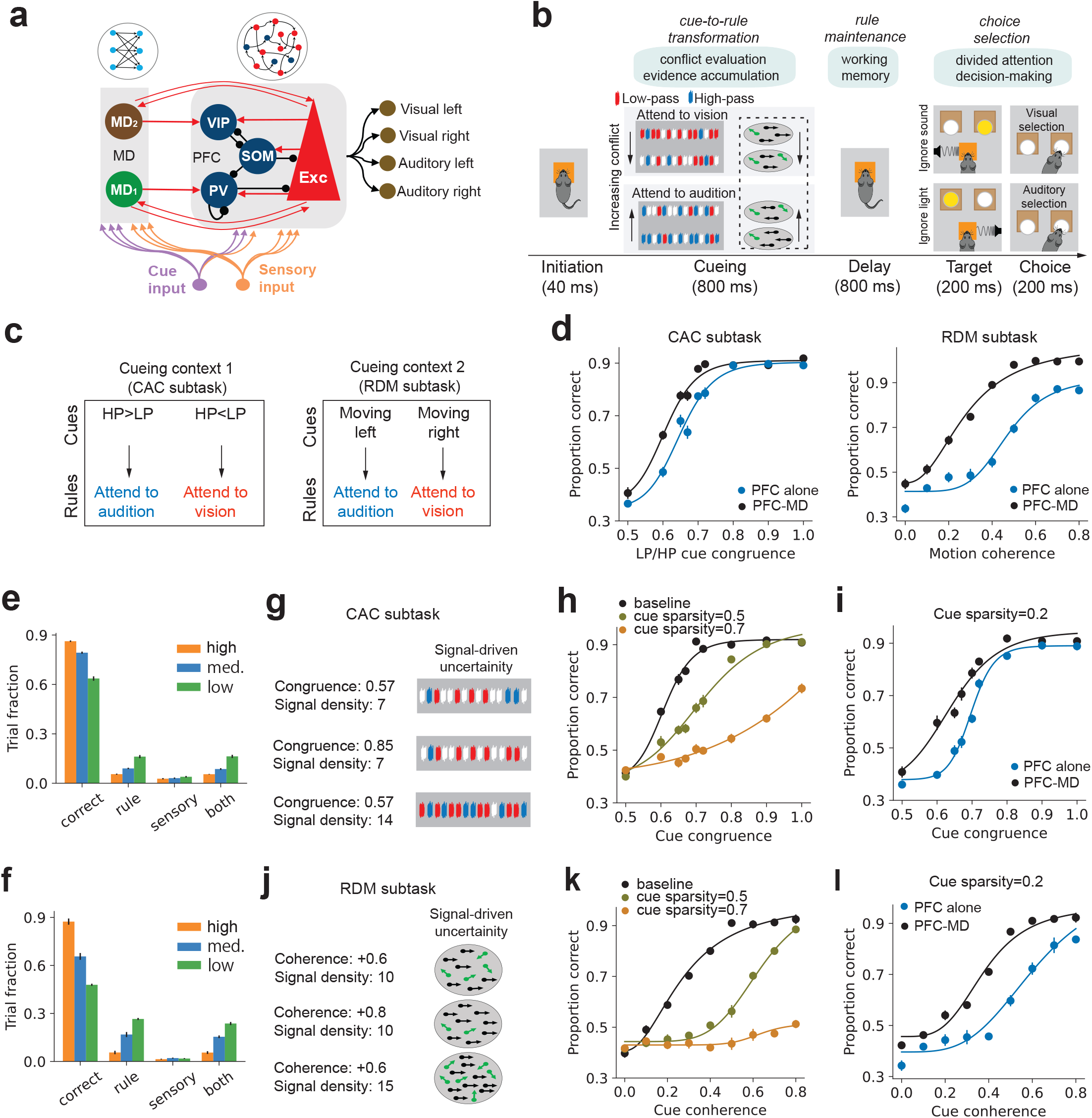
The task-optimized PFC-MD model in decision making with sensory uncertainty. **(a)** Excitation-inhibition (E/I) recurrent neural network (RNN) for modeling the PFC-alone network, where three major prefrontal interneuron cell types were specified. The PFC-MD model consisted of two target-specific non-recurrent excitatory MD subpopulations and bidirectional MD-PFC projections. **(b)** Schematic of a cross-modal, context-dependent decision-making task with working memory and divided attention components. Sensory cues were either conflicting LP/HP pulses or moving random dots. **(c)** Schematic of context-dependent rule encoding. The RDM and CAC subtasks learned the same rule: attend-to-audition vs. attend-to-vision. **(d)** Psychometric curves of RDM and CAC subtasks. Error bar was shown in mean ± s.e.m. Each condition was repeated 10 times with different input realizations yet with identical summary statistics. The PFC-MD model outperformed the PFC-alone model in the presence of intermediate-to-high cue conflict. (**e and f**) Percentage of task error types in two subtasks under various cue uncertainty conditions. **(g)** Illustration of three types of cues with two congruence levels and two densities in the CAC subtask. **(h)** Psychometric curve of the PFC-MD model under three different cue sparsity levels in the CAC subtask. **(i)** Psychometric curve comparison between the PFC-MD and PFC-alone models in the CAC subtask in one cue sparsity condition. **(j)** Illustration of three types of cues with two coherence levels and two densities in the RDM subtask. **(k)** Psychometric curve of the PFC-MD model under three different cue sparsity levels in the RDM subtask. (**l**) Psychometric curve comparison between the PFC-MD and PFC-alone models in the RDM subtask in one cue sparsity condition.

In the first version of context-dependent decision-making task, the task uncertainty contained sensory uncertainty only and two contexts were learned simultaneously. The two contexts corresponded to two cue-independent subtasks that mapped different modalities of sensory cues to one set of rules: attend-to-audition vs. attend-to-vision (**Fig. 1c**). Importantly, the sensory uncertainty was parameterized in the two subtasks: a random dot motion (RDM) subtask and a conflicting acoustic cue (CAC) subtask (see Methods). Additionally, the sparsity of sensory cues, as a generalized form sensory uncertainty, was also parameterized and controlled. These two subtasks consisted of three common task phases: (i) identifying the rule by accumulating sensory evidence from ambiguous cues (800-ms cueing period); (ii) maintaining the rule information during working memory (800-ms task delay period); (iii) making choices in the presence of both visual and auditory stimuli (200-ms target period). Therefore, the model first learned to accumulate sensory evidence of conflicting cues to identify the rule, then learned to preserve the rule information in working memory, and ultimately learned to attend audition/vision correctly and choose a rule-guided target.

We trained PFC-MD models to learn the RDM and CAC subtasks simultaneously where the sensory cueing uncertainty was minimal. Upon reaching the desired task accuracy, we tested the task-optimized PFC-MD models by systematically varying the degree of cue uncertainty. In the RDM context, this corresponded to changing the coherence of random dot motion; whereas in the CAC context, it corresponded to changing the congruence levels of low-pass (LP) vs. high-pass (HP) tones. Namely, the cue-to-rule transformation was determined by motion direction in the RDM context, whereas cue-to-rule transformation was determined by the tone majority.

### PFC-MD model outperformed PFC-alone model in context-dependent decision-making under sensory uncertainty

In testing the model’s generalization, we produced the model’s psychometric curves separately for two subtasks (**Fig. 1d**). By changing the level of coherence or congruence of the cue signals, we found that the PFC-MD model outperformed the PFC-alone model in both contexts, especially in the medium-to-high cue uncertainty ranges. This was consistently observed in multiple independently performance-optimized models (**Supplementary Fig. 1a**), which achieved the same performance on similarly constructed training trials. Furthermore, we examined the percentage of error types in decision making. Given the outcome mismatch among four choices, task errors could be ascribed by either attending the wrong rule (“rule error” during the cueing or delay period) or attending the correct rule but with a wrong choice (“sensory error” during the target period). While testing the task-optimized PFC-MD model, the sensory error was stable regardless of the change in sensory uncertainty, where the rule error changed according to the level of cue ambiguity (**Fig. 1e, f**).

The signal-to-noise ratio (SNR) of sensory cueing was determined by not only the ratio of conflicting cues, but also the density of cue signals (or equivalently the sparsity: the higher the density, the lower the sparsity). For instance, in the CAC context, a 4/2=2:1 ratio of LP/HP pulses indicates a relatively high level of noise---where signals are conflicting and sparse; an identical 8/4=2:1 ratio of LP/HP but with more pulses has a higher SNR, where a higher density of pulses indicates a higher signal level (**Fig. 1g**). Increasing the cue sparsity had a negative impact on the task performance (**Fig. 1h**), but the PFC-MD model consistently outperformed the PFC-alone model (**Fig. 1i** and **Supplementary Fig. 1b**). In the RDM context, we changed the spatial density by varying the number of moving dots with the same cue coherence (**Fig. 1j**). Similarly, decreasing the signal density or increasing cue sparsity led to a degraded task performance (**Fig. 1k**), and the PFC-MD model also outperformed the PFC-alone model (**Fig. 1l** and **Supplementary Fig. 1c**).

### PFC and MD units showed diverse tunings and population dynamics according to task variables, task phase and cue uncertainty

We examined the emergent tuning properties of single units of trained PFC-MD model with respect to task variables during both cueing and delay periods. A majority (∽55-65%) of PFC excitatory units exhibited tuning preference in rule, but not context; some showed cueing context-invariant rule tuning during the delay period (**Fig. 2a, 2d**, and **Supplementary Fig. 2a** for additional examples). Similarly, some (∽60%) of PFC inhibitory units encoded rules during the delay period (**Fig. 2d**). In contrast, ∽80% MD units showed modulation selectivity with respect to the cueing context, but not rule (**Figs. 2b, 2c, 2e** and **Supplementary Fig. 2b** for additional examples). To simulate an effect of evoked firing activity similar to optogenetic MD activation, we induced a transient input to the specific MD subpopulation to increase the phasic firing activity of MD units (Methods). During the baseline (i.e., absence of stimuli), increasing MD_2_ unit firing tended to amplify the firing rate of task-relevant, rule-tuned PFC units, whereas increasing MD_1_ unit firing tended to suppress the firing rate of task-irrelevant PFC units (especially those with lower firing rates) (**Supplementary Fig. 2c**).

**Figure 2.**
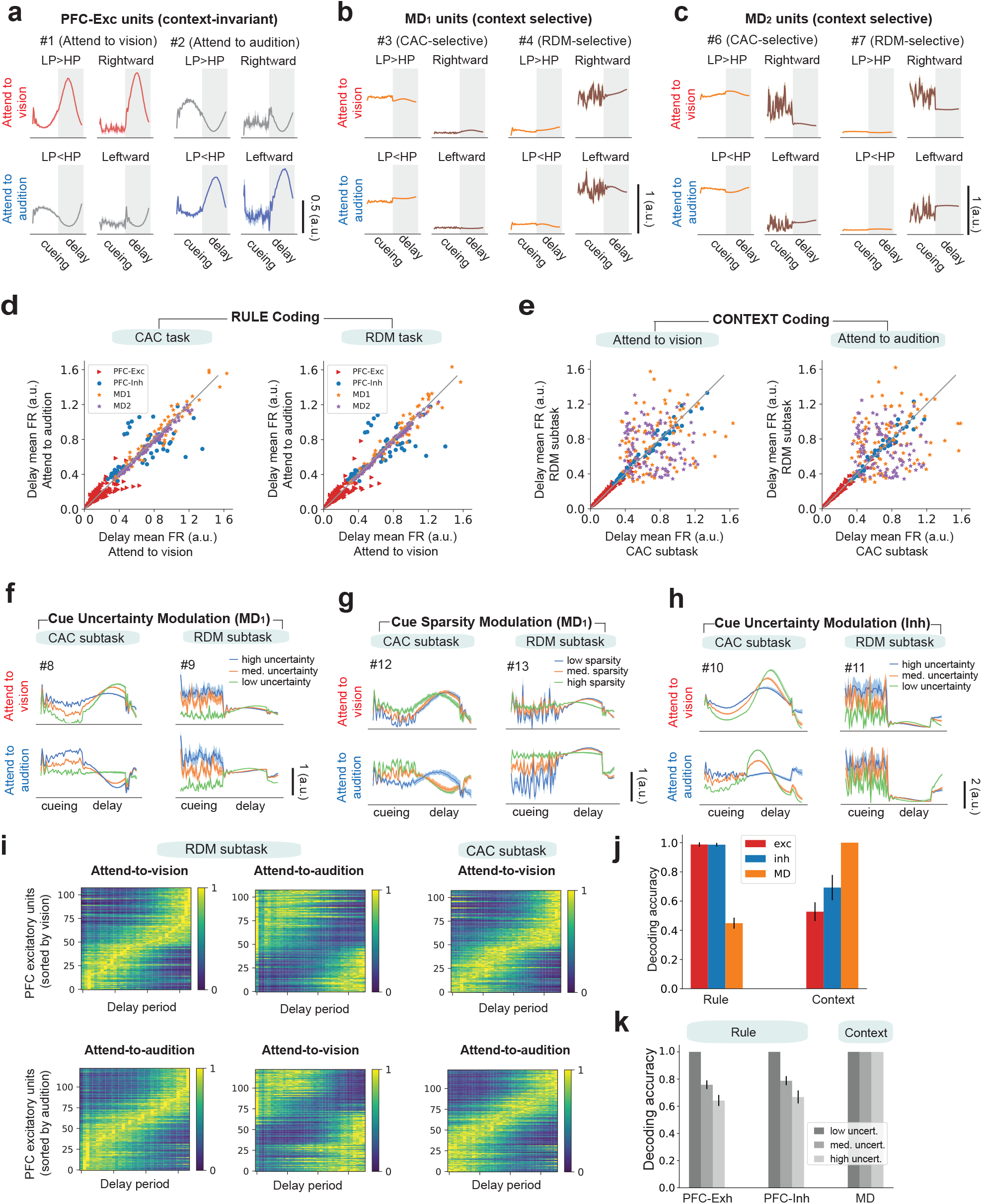
Neural representations of PFC and MD units from the task-optimized PFC-MD network. **(a)** Two representative PFC excitatory units encoding two rules under two contexts. These two units were cue invariant units that encoded the same rule. **(b)** Two representative PFC inhibitory units encoding the cueing context. **(c)**Two representative MD units encoding context but not rule. **(d)** Population statistics of mean firing rates (during the task delay period) of excitatory PFC units and MD units for encoding rule. **(e)** Population statistics of mean firing rate (during the task delay period) of excitatory PFC units and MD units for encoding context. **(f)** Two representative MD_1_ units that showed firing rate modulation with respect to cue uncertainty during both cueing and delay periods. **(g)** The same MD_1_ units also showed firing rate modulation with respect to cue sparsity. **(h)** Two representative PFC inhibitory units that showed firing rate modulation with respect to cue uncertainty during both cueing and delay periods. **(i)** Prefrontal neural sequences showed rule specificity and context invariance. Each heatmap shows the normalized peri-stimulus time histogram (PSTH) of selected prefrontal excitatory units of the task-optimized PFC-MD model during the delay period. In the first row, all units of all panels were sorted by the same order according to attend-to-vision tuning. In the second row, all units of all panels were sorted by the same order according to attend-to-audition tuning. The first and second columns demonstrated rule specificity, whereas the first and third columns demonstrated context invariance. **(j)** Population decoding analysis showed that the PFC population better encoded rule, whereas the MD population better encoded context. Error bar denotes s.e.m. (n=20). **(k)** Increased in cue uncertainty caused decreased rule decoding accuracy for the PFC, but did not affect context decoding accuracy for the MD. Error bar denotes SD (n=20). In rule decoding, all paired comparisons were statistically significant (P<7×10^-8^, two-tailed Wilcoxon rank-sum tests).

Additionally, during the cueing period, some MD_1_ units (25-30%, across subtasks and multiple test conditions) modulated their firing rates with respect to cue uncertainty (**Fig. 2f** and **Supplementary Fig. 2d**). We also found firing modulation with respect to cue sparsity for the MD_1_ subpopulation (**Fig. 2g**), but not for MD_2_ units (**Supplementary Fig. 2e**). Among MD_1_ units, tunings with respect to context and cue uncertainty were relatively orthogonal since very few MD_1_ units showed conjunctive coding (**Supplementary Fig. 2f**). Similar cue uncertainty tunings were also found in PFC inhibitory units (**Fig. 2h**). It is noteworthy that not only these emergent PFC and MD tuning properties were in line with experimental findings in task-performing mice^5,6^, intensive computer simulations also enabled us to investigate the dependency of these tunings with respect to thalamocortical connectivity, providing insight into “why” and “how” questions of MD regulation (see Discussion).

Increasing cue uncertainty or cue sparsity reduced rule selectivity of PFC excitatory units in working memory (**Supplementary Fig. 3**), suggesting that reducing the SNR of sensory cues may lead to an information loss during cue-to-rule transformation. During the delay period, sorting the peak responses of rule-tuned PFC excitatory units produced a rule-specific sequence^17,24,26,28^; moreover, the neural sequence was context-invariant for RDM and CAC subtasks (**Fig. 2i**). To further explore the tuning properties of PFC-MD populations, we employed population decoding analyses to read out either rule or context information on a trial-by-trial basis based on the mean firing of population (Methods). The rule information could be reliably decoded from the PFC, but not from the MD (**Fig. 2j**). Increasing cue uncertainty led to a decrease in the PFC’s rule-decoding accuracy, but had no effect on the MD’s context-decoding accuracy (**Fig. 2k**).

Furthermore, dimensionality reduction of the PFC population activity revealed low-dimensional neural trajectories in the task-specific subspace (**Fig. 3a, d**). Increasing the level of cue uncertainty or sparsity led to changes of neural trajectories (**Fig. 3b, c, e, f**). Interestingly, the neural trajectory converged to fixed points faster under the lowest sensory uncertainty, and the initial velocity was greater while dealing with lower cue uncertainty (**Fig. 3g, h**). All neural trajectories reached to a steady state where the velocity was close to zero. A fixed-point analysis on the three-dimensional PC subspace revealed two “line attractor-like” basins, with each one matching a rule (attend-to-vision vs. attend-to-audition) and aligning sensory uncertainty from one end to the other (**Fig. 3i**). Collapsing these points could reveal two distinct line attractor-like profiles with finite line length; the neural trajectory could diverge from the attractor basins when the sensory cue uncertainty was far too high.

**Figure 3.**
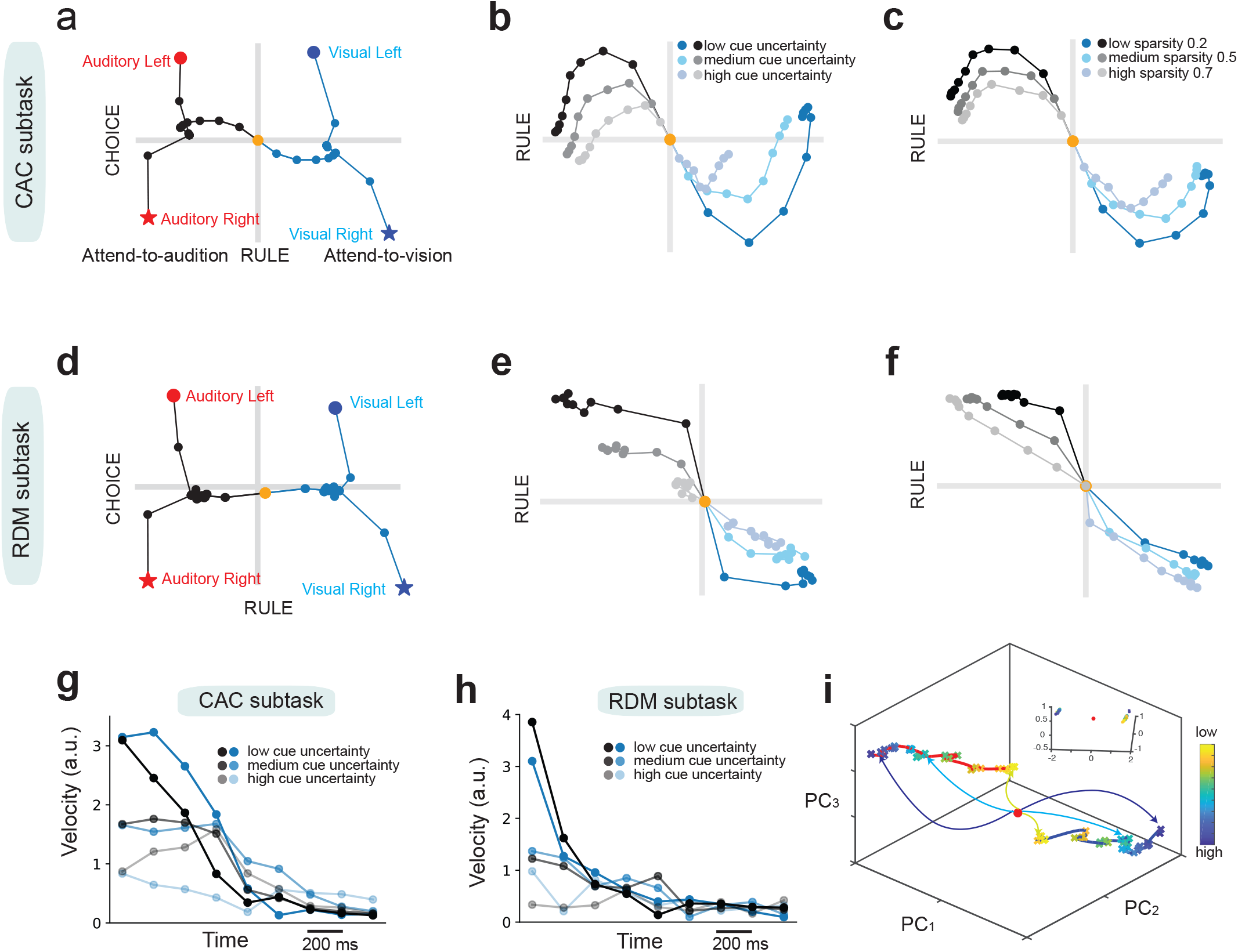
Population representations from the task-optimized PFC-MD model. **(a)** Dynamics of population responses in the CAC subtask. The average population trajectory for a given condition and time was represented as a point in the state space. Responses from correct trials only were shown from the cue onset to the end of target period (80-ms step size), and were projected into the two-dimensional subspace capturing the variance according to the rule (attend-to-audition vs. attend-to-vision) and choice (see Methods). Units are arbitrary. The origin represents the cue onset. **(b)** Similar to (**a**), except only during the cueing period. For a better illustration, three levels of cue uncertainty were shown by different shades of grey or blue color (dark/intermediate/light shade: low/medium/high cue uncertainty, respectively). **(c)** Similar to (**b**), except for a fixed cue uncertainty but different levels of cue sparsity. Three levels of sparsity were shown by different shades of grey or blue color (dark/intermediate/light shades: low/medium/high sparsity, respectively). (**d-f**) Similar to (**a-c**) but for the RDM subtask. (**g and h**) Comparison of neural velocity during both cueing and delay periods for different levels of cue uncertainty for the CAC and RDM subtasks, respectively. The change of neural activity reduced to a low level during the delay period, reaching a fixed-point regime. (**i**) Fixed-point analysis in the three-dimensional PCA subspace revealed two “line-attractor-like” basins, with each basin representing a rule. Each cross symbol corresponded to a fixed point (color sorted by cue uncertainty). The red origin represents the cue onset. Inset: rotating the view angle and collapsing these points revealed two “fixed-point-like” attractors.

### MD enhances working memory maintenance in the PFC-MD network

The neural basis for persistent activity in working memory is thought to involve recurrent excitation among prefrontal excitatory neurons. During working memory, the rule information is maintained in thalamocortical communications, but the information may be subject to a loss with increased task delay duration or noise interference. Therefore, we tested the robustness of task-optimized PFC-MD model subject to a potential information loss. In so doing, we gradually increased the duration of delay period from the initial 800 ms to 1200 ms, and found that increasing delay duration had a negative impact on task performance (**Fig. 4a**; similarly for PFC-alone network, **Supplementary Fig. 4a**). To demonstrate the role of MD in working memory maintenance,^17^ we increased the firing rate of MD_1_ or MD_2_ subpopulation during an elongated 1100-ms delay period (with coherence/congruence of 0.8 in respective RDM/CAC subtasks), which led to a slight boost in rule maintenance. Notably, it required less firing rate elevation for MD_2_ than MD_1_ to achieve the same task improvement (**Fig. 4b, c**), probably because the MD_2_⟶VIP pathway has an amplification effect on increasing the target sensitivity for working memory maintenance. In contrast, reducing the MD_1_ activity during the delay period had an opposite effect on the task performance (**Supplementary Fig. 4b**). Alternatively, we strengthened or weakened prefrontal Exc-to-MD corticothalamic connection strengths and observed similar effects on performance (**Supplementary Fig. 4c-f**). Together, these results support that reciprocal PFC-MD communications help maintain the rule information through two functionally distinct thalamocortical//corticothalamic pathways.

**Figure 4.**
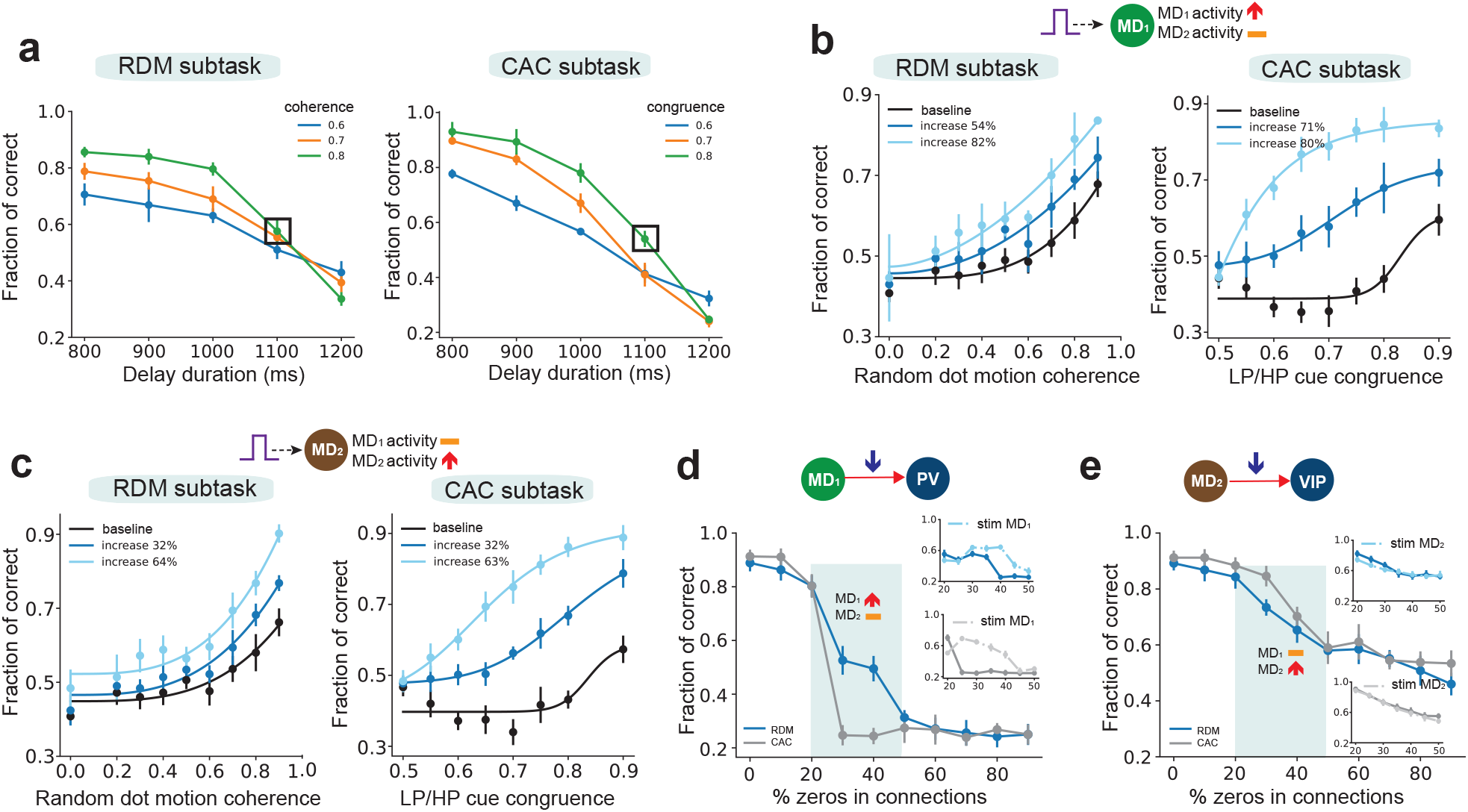
MD enhances working memory maintenance in the PFC-MD network. **(a)** Increasing the duration of task delay period reduced the task performance under cue uncertainty in both subtasks. Figure legend denotes the level of cue coherence or congruence. Error bar denotes SD (n=10). The task condition marked with “ ” was used in the illustrations of three remaining panels. (**b and c**) Increasing the MD_1_ or MD_2_ population firing rate during an elongated delay period improved the working memory and the psychometric curve in both subtasks. Black curve denotes the baseline, and the number of % denotes the relative increase in subpopulation firing rate. **(d)** Weakening MD_1_⟶PV connections (by increasing the percentage of zeros) during the cueing period quickly reduced the task performance in both RDM and CAC subtasks. Inset: increasing MD_1_ activity could rescue each subtask performance under a wide range of connectivity conditions (shaded area: 20%-50% of zeros). Error bar denotes SD (n=10). **(e)** Weakening MD_2_⟶PV connections only degraded the task performance slowly in both RDM and CAC subtasks, but increasing MD_2_ activity had no effect on the change in task performance (see inset).

To distinguish the role of two specific thalamocortical projections (MD_1_⟶PV and MD_2_⟶VIP) in regulating prefrontal computation, we disconnected one of two thalamocortical projections (i.e., “computational lesion”) while keeping the other intact. We found that weakening MD_1_⟶PV connectivity (by setting a small percentage of connections to zeros) reduced the performance rapidly for RDM and CAC subtasks (**Fig. 4d**). Enhancement of phasic MD_1_ activity could boost task performance, but only within a narrow range (**Fig. 4d**, dashed line within the inset). In contrast, weakening MD_2_⟶VIP connectivity caused task performance to decay slowly (**Fig. 4e**); however, enhancement of MD_2_ activity had little effect on task performance (see inset). This was possibly because there was no direct VIP⟶Exc pathway to prefrontal excitatory neurons.

### Probing mechanistic causes of the PFC-MD circuit in cognitive deficits

Our task-optimized PFC-MD models may serve as a platform to interrogate mechanisms of abnormal neural representations and cognitive deficits in working memory or behavioral inflexibility. This was achieved by modifying the “normal control” with specific “computational lesion or dysfunction”.

Recurrent synaptic excitation among pyramidal neurons is approximately balanced by synaptic inhibition^32^. Thus, we first changed the overall inhibition to excitatory neurons and induced cortical excitation-inhibition (E/I) imbalance; this was done by increasing the sparsity of prefrontal SOM⟶Exc connections to reduce inhibition. Alternatively, MD inhibition also alters prefrontal E/I balance^33^. Notably, these operations changed the rule tunings of some PFC excitatory units during the delay period and reduced the choice discriminability at the population level during the target period (**Fig. 5a, b** and **Supplementary Fig. 5a, b**). This can be explained by the fact that the prefrontal network dynamics became more unstable and sensitive to external input perturbation, which pushed neural trajectories out of the stable attractor space. Visualizing population dynamics in the PCA subspace showed that E/I imbalance distorted neural trajectories (**Fig. 5c** and **Supplementary Fig. 5a, b**), so that the network might favor one rule over the other (i.e., “cognitive biases” observed in many mental disorders).

**Figure 5.**
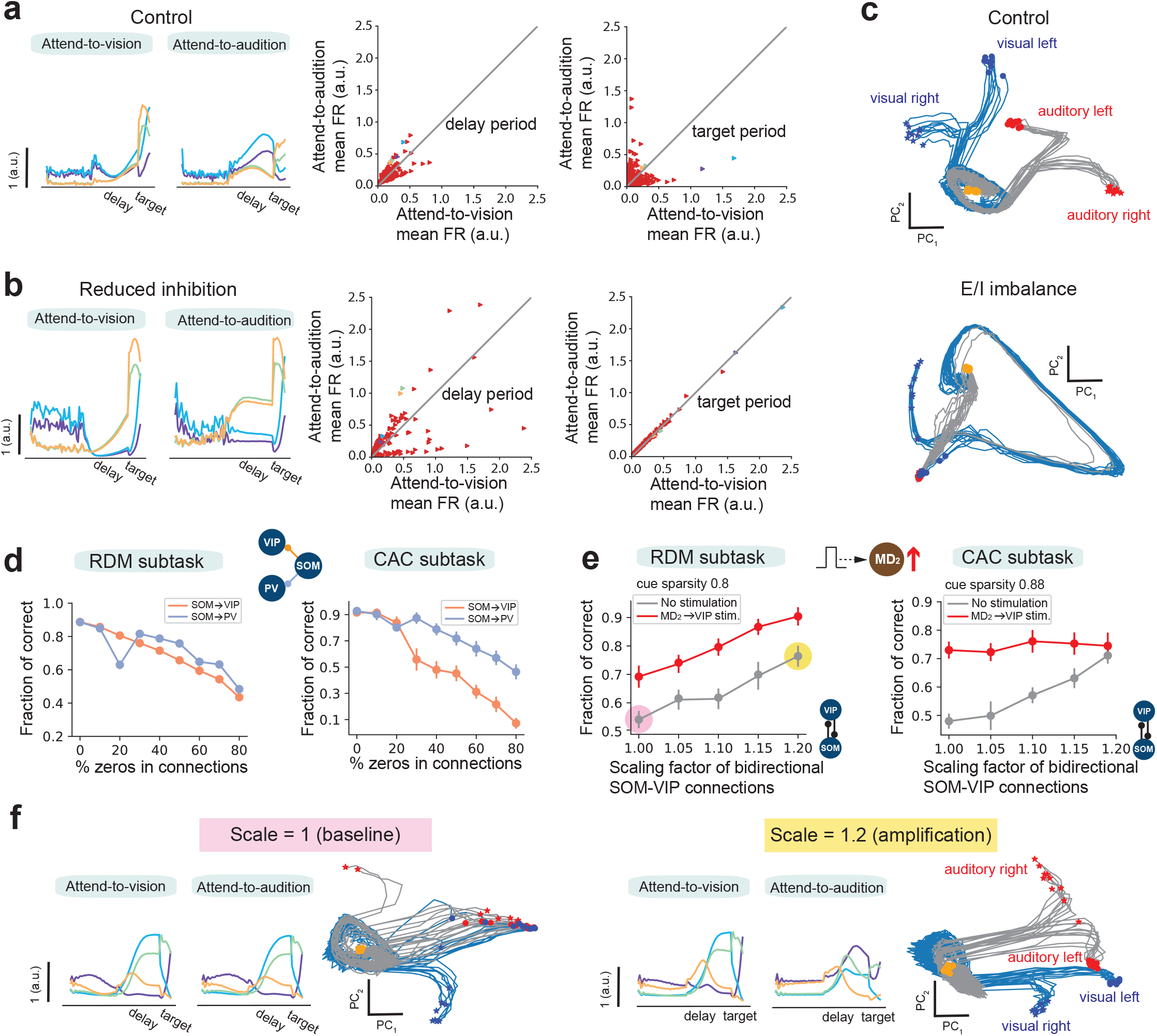
Parsing cognitive deficits and probing mechanistic causes in the PFC-MD networks. **(A)** During the control condition, PFC excitatory units (4 selected units are indicated by different colors) preserved rule tunings in both delay and target periods (left panel). Mean firing rate (FR) comparison of PFC excitatory units between two rules in delay (middle panel) and target (right panel) periods. Four units in the left panels are labeled in the same color in both middle and right panels. **(B)** In the presence of E/I imbalance (e.g., reduced inhibition), PFC excitatory units shown in (**A**) increased their firing rates but decreased their rule discriminability during the target period. **(C)** Two-dimensional neural trajectory representations in control and E/I imbalance conditions. Trajectories were generated from all correct and error trials. Orange dots: cue off and start of delay period. Blue and red end points represent two rule representations, whereas dot and star symbols represent left and right choices, respectively. **(D)** The task performance decreased with reduced prefrontal SOM⟶VIP and SOM⟶PV connectivity. **(E)** Increasing the mutual inhibition strength between SOM and VIP neurons amplified the gain under a high cue sparsity. Stimulating the MD_2_⟶VIP pathway further facilitated the amplification and improved the task performance. Two shaded circles indicate the conditions illustrated in (**F**). **(F)** In the case of RDM task of (**E**) (with cue sparsity 0.8), comparison of PFC excitatory unit tunings and two-dimensional neural trajectories between scale=1 (light gray, baseline) and scale = 1.2 for bidirectional SOM-VIP connection strengths. In the latter case, single-unit rule tunings emerged and population responses improved rule discriminability.

We further weakened prefrontal SOM⟶VIP and SOM⟶PV connectivity independently and observed a more robust task performance in SOM⟶PV manipulations (**Fig. 5d**), while a similar manipulation of VIP⟶SOM connectivity had little impact (**Supplementary Fig. 5c**). For bidirectional SOM-VIP connections, strengthening mutual inhibition between SOM and VIP neurons could amplify the sparse cue signal (grey curves in **Fig. 5e**; note that this was effective for bidirectional SOM-VIP amplification but not for unidirectional manipulation; **Supplementary Fig. 5d**), supporting the role of VIP-SOM motif in gain control^19^. A closer examination of single-unit and population responses revealed that bidirectional SOM-VIP amplification produced emergent rule tunings and enhanced rule discriminability (**Fig. 5f**). Additional MD_2_⟶VIP stimulations further facilitated this SOM-VIP amplification (red curves in **Fig**. 5e), validating the concept of MD amplifier to regulate prefrontal computation^5^. In animal experiments, abnormal cortical GABAergic signaling or deficits in interneuron subtype has been implicated in cognitive impairment in autism spectral disorder (ASD), such as decreased responses to salient stimuli (“hypo-sensitivity”) under a low SNR or decreased inhibitory gain control^34,35^.

### Unbalanced corticothalamic connectivity reshapes MD tunings

Next, we investigated whether modified corticothalamic connectivity could change MD neuronal tunings. Data from previous mouse experiments have suggested that MD_GRIK4_ neurons received dense inputs from L5/6 of the prelimbic cortex, in contrast to MD_DRD2_ neurons that receive sparse prefrontal inputs (experimental data not shown). To incorporate this prior knowledge, we made two modifications to the PFC-MD model. First, we assumed unbalanced MD_1_ and MD_2_ subpopulations (3:2 ratio in place of the previous 1:1 ratio). Second, we assumed that MD_2_ units received a much sparser projection from prefrontal excitatory units than MD_1_ units and imposed a sparsity constraint during the course of model training, where Exc⟶MD_2_ connectivity was 75% sparser than Exc⟶MD_1_ connectivity (**Fig. 6a**). Notably, this modified corticothalamic connectivity did not change psychometric curves of the task-optimized PFC-MD model (**Fig. 6b**), but led to emergent weak tunings in some MD_2_ units with respect to sensory uncertainty (**Fig. 6c**). Additionally, MD_1_ stimulation, but not MD_2_ stimulation, improved the performance of two subtasks under cue uncertainty; yet either MD_2_ or MD_1_ stimulation could improve the task performance under cue sparsity (**Fig. 6d, e**). Furthermore, when cue uncertainty and sparsity were both present, stimulation of both MD subpopulations was required in order to improve task performance (**Fig. 6f**). Together, these findings further suggest distinct modulatory functions imposed by cell type-specific connectivity-dependent thalamocortical projections.

**Figure 6.**
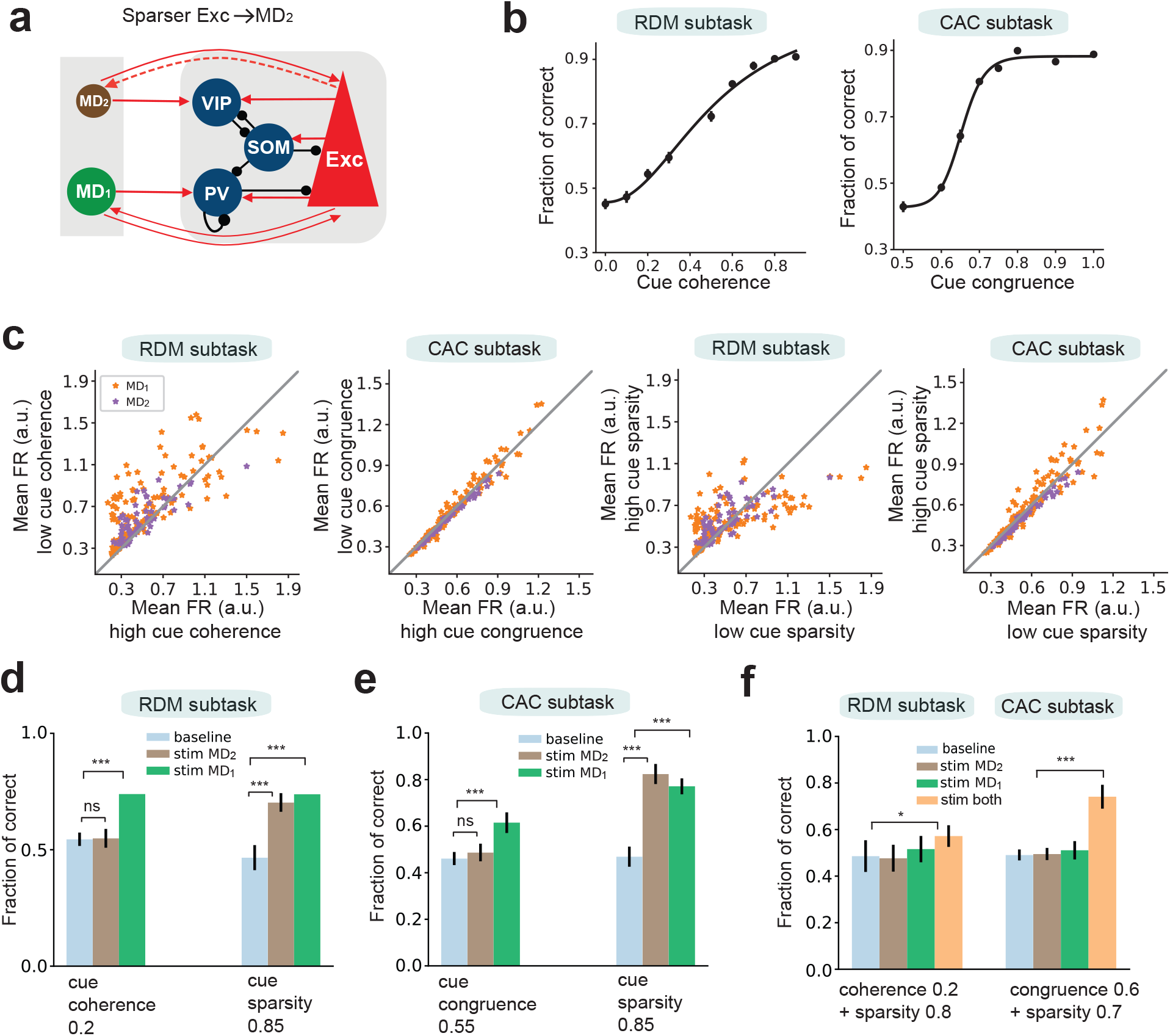
Modified PFC-MD model with additional sparse corticothalamic connectivity. **(a)** The modified PFC-MD model that assumed unbalanced MD subpopulations (as shown by two different circle sizes) and sparser Exc⟶MD_2_ connectivity than Exc⟶MD_1_ connectivity (as shown by a thinner connection). **(b)** The psychometric curves of task-optimized modified PFC-MD model in two subtasks. Error bar was shown in mean ± s.e.m. (n=10 realizations). **(c)** Population statistics of mean firing rates (during the cue period) of MD_1_ and MD_2_ units for encoding cue uncertainty and cue sparsity. (**d and e**) MD_1_ stimulation, but not MD_2_ stimulation, improved task performance under cue uncertainty. MD_1_ or MD_2_ stimulation improved task performance under cue sparsity. Error bar denotes SD (n=10). Cue uncertainty comparison during the RDM (CAC) subtask: baseline vs. MD_2_ stimulation, P=0.82 (0.16); baseline vs. MD_1_ stimulation, P=0.00016 (0.00016); MD_2_ vs. MD_1_ stimulation, P=0.00016 (0.00043). Cue sparsity comparison during the RDM (CAC) subtask: baseline vs. MD_2_ stimulation, P=0.00016 (0.00016); baseline vs. MD_1_ stimulation, P=0.00016 (0.00016); MD_2_ vs. MD_1_ stimulation, P=0.045 (0.017), two-tailed Wilcoxon rank-sum tests. (**f**) When both cue uncertainty and cue sparsity were present, MD_1_ or MD_2_ stimulation alone couldn’t improve task performance, but simulation of both MD subpopulations could. Baseline vs. stimulation both,*, P=0.011; ***, P= 0.00015, two-tailed Wilcoxon rank-sum tests. Other paired comparisons were n.s.

### Bidirectional PFC-MD synaptic plasticity enabled rapid learning of context switching

In the second version of context-dependent decision-making task, both sensory and mapping uncertainties were present and two contexts were learned sequentially. Specifically, we introduced cue-to-rule mapping uncertainty and sequential context switching (*Context 1*⟶*Context 2*⟶ *Context 1’*) using a modified version of CAC subtask (**Fig. 7a** and Methods). Upon learning Context 1, we assumed a prewired intracortical structure and kept the synaptic connectivity intact within the PFC network. Interestingly, PFC inhibitory-to-excitatory connectivity formed two distinct patterns according to the rule tunings of excitatory units (**Supplementary Fig. 6a**); further examination of all presynaptic connections to postsynaptic PFC excitatory units revealed distinct clusters according to the rule-tuning profiles (**Supplementary Fig. 6b**).

**Figure 7.**
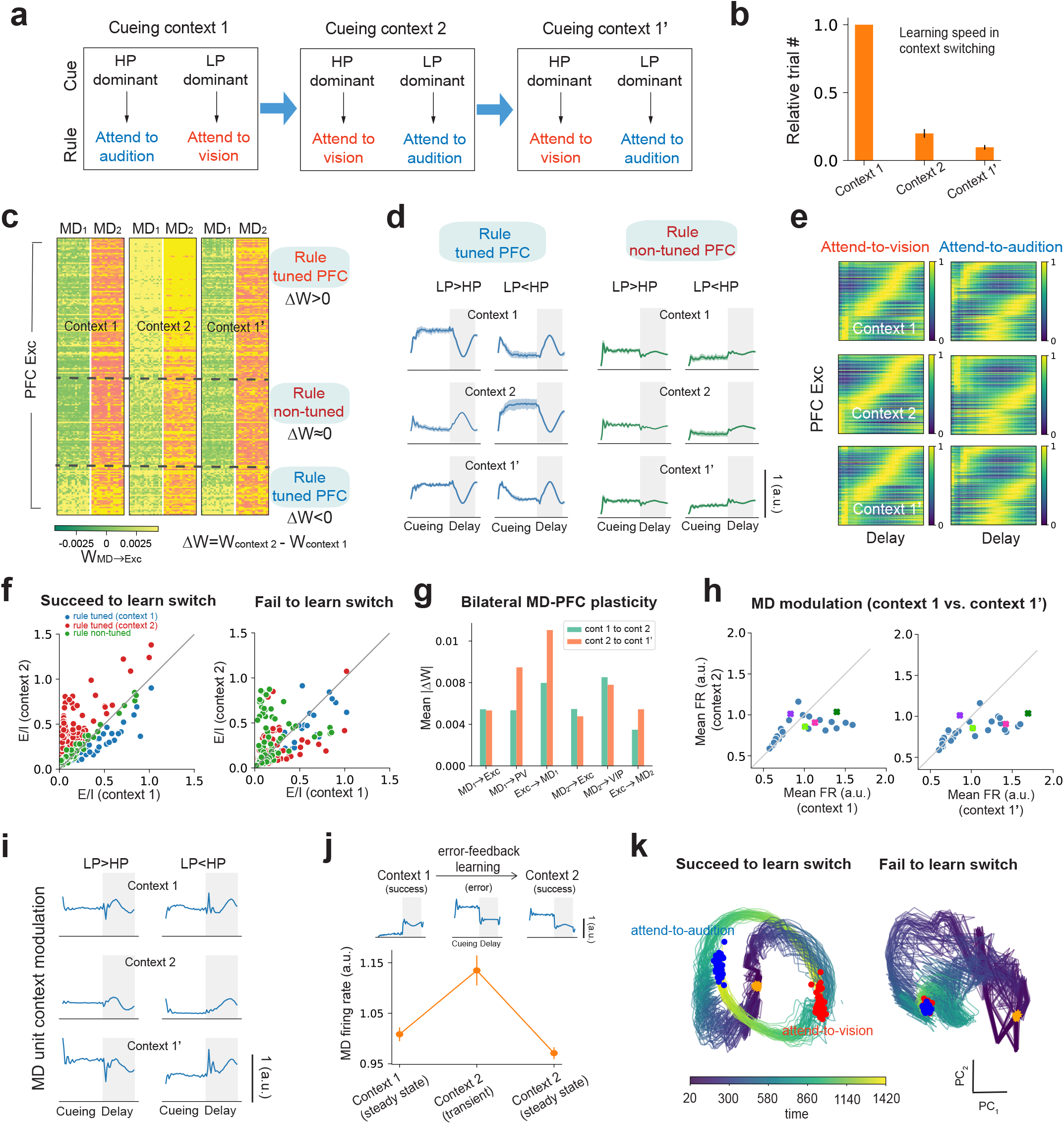
Thalamocortical plasticity in the PFC-MD model enabled context switching to tackle mapping uncertainty. **(a)** Schematic diagram of the context-switching task (Context 1⟶Context 2⟶Context 1’). **(b)** Relative learning speed in context switching, where neural plasticity of local MD-PFC connectivity was induced in trial-by-trial learning of Context 2 and Context 1’. Error bar denotes s.e.m. (n=10). **(c)** Heatmap of connectivity of MD (MD_1_ and MD_2_) to prefrontal excitatory (Exc) units. Let W denote the MD-to-Exc connection matrix, units were sorted based on the change of ΔW=W_context 2_-W_context 1_. According to the synaptic change ΔW, PFC excitatory units were mapped to two functional cell types: rule tuned (red and blue) vs. non-tuned (green) units. **(d)** Tuning curve examples of rule tuned (blue) and non-tuned (green) PFC units. Shaded area denotes the task delay period. **(e)** PFC excitatory units showed context-invariant rule-specific sequential activity during the delay period. Each row of the heatmap corresponded to the normalized trial-averaged firing activity. Units were sorted based on the location of peak firing rates and ranked in the same order in all six panels. **(f)** The E/I input of PFC excitatory units during the task delay period were clustered according to the PFC rule tuning properties when the PFC-MD network succeeded to learn the context switch (left). The cluster structure was lost when the PFC-MD network failed to learn the context switch (right). Units are color coded using the same color scheme based on rule-tuned (red and blue) or non-tuned (green) property in (**C**). **(g)** Quantification of average plasticity in bilateral MD-PFC connections during context switching. **(h)** MD units showed context-invariant firing. Note that some MD units (overlaid “×” symbols) preserved their mean firing rates during the delay period between Context 1 and Context 1’. **(i)** Turning curve illustrations of one MD unit during context switching. Shaded area denotes the task delay period. Notice the strong context modulation between different contexts, but little modulation with respect to the rule. **(j)** A subset of MD units (n=8 out of 30 from one trained PFC-MD model) showed increased modulation with respect to decision error during the transient switching stage. The MD firing rate was averaged across the delay period where the change of network state was very small (i.e., steady state). Tuning curves of one MD were shown on the top. **(k)** Comparison of neural trajectories of PFC population dynamics during cueing and delay periods between conditions where the PFC-MD succeed and failed to learn the context switch. Trajectories were color coded to represent time from the cue onset to the end of delay.

Further, we adapted the MD⟶PFC and PFC⟶MD synaptic connections based on inter-trial error feedback to learn context switching in a sequential manner (i.e., on a trial-by-trial basis). Notably, modification of thalamocortical and corticothalamic connections allowed the pretrained PFC-MD network to learn the new cue-to-rule transformation rapidly (**Fig. 7b**). We found that MD⟶Exc pathways played a critical role in context switching and rule remapping (**Fig. 7c**). The change in thalamocortical connections also mapped to functional cell types in PFC excitatory units, majority of which (>60%) displayed rule-invariant tunings despite context switching (**Fig. 7d**). Some PFC excitatory units did not change firing rates but shifted their peak firing rates temporally during the delay period (**Supplementary Fig. 6c**). Together, the PFC excitatory population displayed robust rule-invariant sequence during the delay period (**Fig. 7e**). Furthermore, the rule-tuning of PFC excitatory units tended to cluster with specific E/I ratio and modified the E/I input accordingly after successful context switching (**Fig. 7f**, left panel); in contrast, the PFC tuning properties would fail to discriminate the rule if context switching was unsuccessful (**Fig. 7f**, right panel).

On the other hand, MD units showed context-invariant tuning between *Context 1* and *Context 1’* (**Fig. 7h**; and a unit tuning example in **Fig. 7i**). Additionally, the cue-to-rule transformation of the RNN at the steady state are qualitatively similar between *Context 1* and *Context 1’* (**Supplementary Fig 7d** and Methods). Furthermore, a small subset of MD units increased firing with inter-trial error during the course of context-switch learning: they increased the delay-period firing rates during the transient remapping state and resumed the baseline firing when successful context switching was completed (**Fig. 7j**; top: single-unit example; bottom: mean firing rate statistics, n=8 from one trained PFC-MD network). Imbalanced E/I or insufficient MD-to-PFC modulation led to a failure in cue-to-rule remapping at both single-unit and population representations (**Fig. 7f, 7k**).

### Theoretical insight on MD feedforward control

Our biologically-constrained computational models have provided a paradigm to test the role of MD in regulating sensory uncertainty in context-dependent decision-making tasks (**Fig. 8a**). The modeling framework also allows us to compare functions and computational efficiency derived from different intracortical, intra-thalamic, thalamocortical and corticothalamic connectivity (**Fig. 8b**). Bilateral interactions between the PFC and MD are critical for cognitive flexibility^6^. A key question is why the combination of a recurrent E/I structure (PFC) and a feedforward excitatory architecture (MD) plus neuron subtype specificity produces more flexible control and robustness in decision making with task uncertainties, as compared to a PFC-alone model that achieves nearly the same computational power? To further probe this question, we systematically varied the size of MD population (*N*_MD_=2,8,16,30), retrained the PFC-MD network, and then investigated the impact of MD on the learning speed of context switching and network properties. We found that adding more MD units improved the relative switch speed in trial-by-trial learning (**Fig. 8c**). Therefore, neurobiology may teach us a good reverse-engineering lesson. One specific theoretical insight is that the MD may help “slow down” prefrontal dynamics by increasing the time constant in order to improve information integration during cueing or working memory (**Fig. 8d** and Methods). Additionally, our computer simulations suggest that the feedforward MD structure has a lower intrinsic dimensionality than the recurrent PFC structure in the full-size PFC-MD network (**Fig. 8e**).

**Figure 8.**
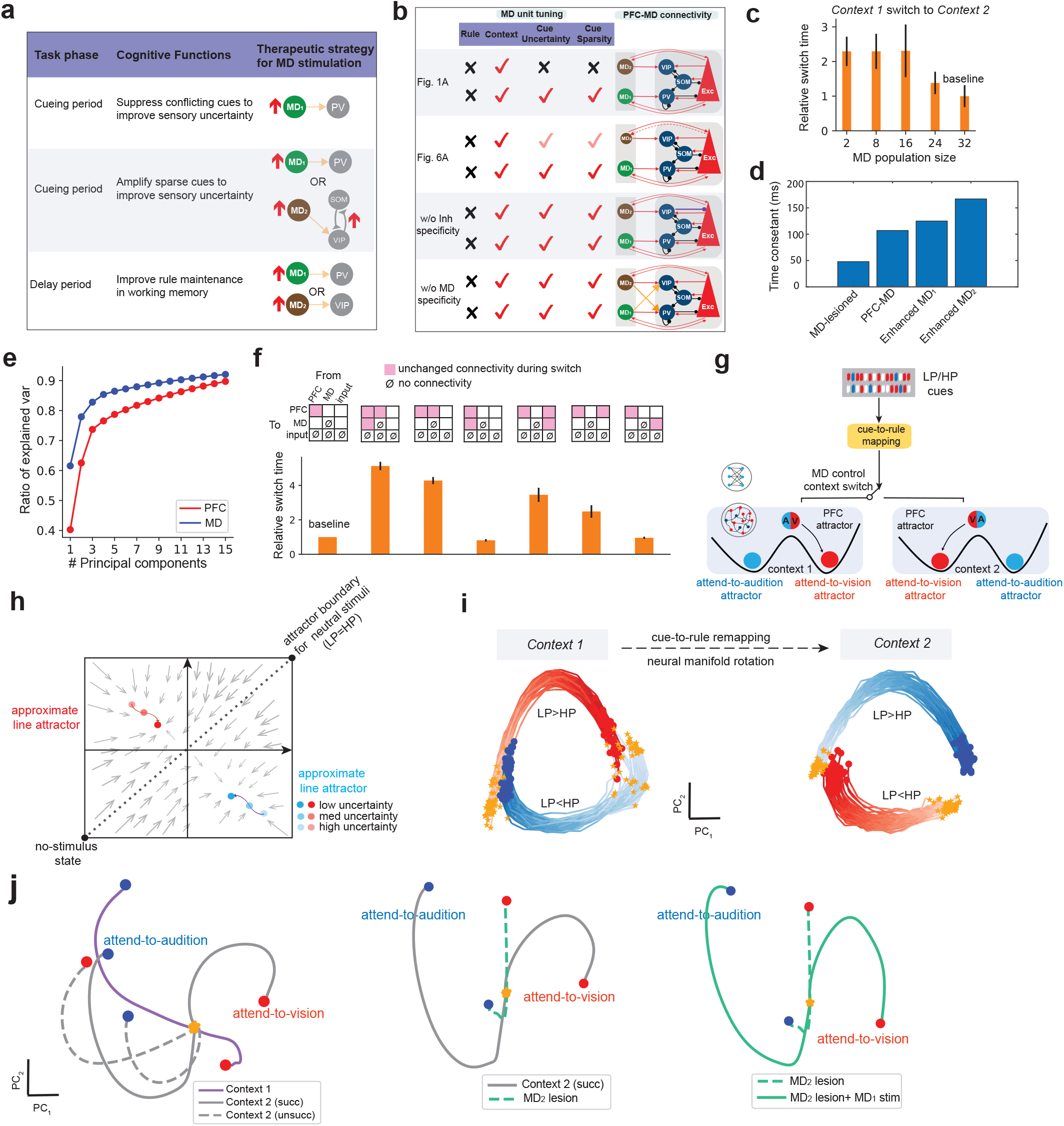
Theoretic insight from computational modeling of the PFC-MD circuit in context-dependent decision making. **(a)** Schematic summary of cell type and task-phase specific regulatory roles of MD thalamus in decision making under sensory uncertainty. **(b)** Differential assumptions of intracortical, thalamocortical and corticothalamic connectivity had a direct impact on the MD unit tuning in task-optimized PFC-MD models. **(c)** Comparison of relative context switch time with different sizes of MD population. All time was normalized with respect to the baseline (MD size of 32). Error bar denotes s.e.m. (n=10). **(d)** Ratio of explained variance of PFC and MD populations derived from the task-optimized PFC-MD model. Our computer simulations and principal component analysis (PCA) showed that the recurrent PFC structure had a larger intrinsic dimensionality than the feedforward MD structure. **(e)** Estimated time constant from the nonlinear dynamical systems under different thalamocortical connectivity manipulations. These results suggest that the MD could “slow down” prefrontal dynamics by increasing the time constant for improving information integration during cueing or working memory period. The MD_1_ and MD_2_ enhancement corresponded to the experiments where respective thalamocortical connections were strengthened during working memory. **(f)** Comparison of relative context switch time with different assumptions of modifiable connectivity during switch. In the top, unilateral or bilateral PFC, MD and input connectivity is shown (∅ denotes no connectivity). In total, there are five unidirectional connections that can be adapted. In the bottom, all time in the y-axis was normalized with respect to the baseline (unchanged intracortical connectivity). Error bar denotes s.e.m. (n=10). **(g)** Schematic illustration of the MD’s role in context switch during cue-to-rule remapping, and the PFC is illustrated as a bistable attractor. **(h)** Two-dimensional (2D) vector field that illustrates the bistable PFC dynamics as an approximate line attractor. The diagonal dotted line represents an attractor boundary for neutral stimuli (e.g., LP=HP). Rotating the 2D vector field by 180 degrees is equivalent to switching the cue-to-rule transformation while preserving the PFC dynamics. **(i)** Comparison of PFC population dynamics under *Context 1* and *Context 2* while projecting them onto the same PCA subspace. The neural trajectories were nearly orthogonal to each other. Each color trace represents a single trial with specific cue input (LP>HP or LP<HP). Light to dark color represents the time evolution of the trial. **(j)** Left panel: Comparison of neural trajectories of PFC dynamics in *Context 1, Context 2* with successful context switching, and *Context 2* with unsuccessful context switching. Middle panel: Comparison of neural trajectories of PFC dynamics in *Context 2* with successful context switching and *Context 2* with MD_2_ lesion. Right panel: Comparison of neural trajectories of PFC dynamics in *Context 2* with MD_2_ lesion and *Context 2* with MD_2_ lesion plus MD_1_ stimulation.

Our model provides new insight into the role of MD-PFC connectivity in enabling rapid context switching to learn a cue-to-rule remapping, and further produces new experimentally testable hypotheses. As control experiments, in addition to keeping intracortical PFC-to-PFC connectivity intact, we systematically kept the unilateral or bilateral MD-PFC synaptic connectivity unchanged during context switching and only allowed synaptic plasticity among the remaining connectivity (**Fig. 8f**, top panels). Our result showed that the PFC rule invariance property was lost when MD plasticity was disrupted, and the speed of cue-to-rule remapping was the slowest when bilateral MD-PFC plasticity was disabled (**Fig. 8f**; compared to the baseline where intracortical connectivity was intact). In contrast, enabling MD⟶PFC plasticity significantly improved the switch speed, suggesting that the crucial role of thalamocortical input in regulating recurrent prefrontal computation for cognitive flexibility. Based on these results, we envisioned that the MD acts like an ON/OFF switch that modulates the respective E/I input of rule-tuned PFC units and changes prefrontal tunings according to the context. If the recurrent prefrontal circuit is viewed as an attractor network (**Fig. 3i**), the context-encoding MD can play the role of controller to usher the PFC to a proper attractor space, facilitating noise reduction (for ambiguous cues) and cue-to-rule remapping (for the same cues) (**Fig. 8g**). If the PFC dynamics is simplified as a two-dimensional (2D) system, we envisioned that the lower-dimensional (i.e., 1D) MD is acting like an actuator on the 2D system; rotating the system by 180 degrees through MD-PFC plasticity may enable cue-to-rule remapping while keeping the original PFC dynamics intact (**Fig. 8h**). This intuition was confirmed by our computer simulations, where prefrontal population dynamics under *Context 1* and *Context 2* were projected onto the same subspace (**Fig. 8i**). Furthermore, the orthogonal relationship of PFC population dynamics was lost if the PFC-MD network failed to learn the context (**Fig. 8j**, left panel, dashed lines). Additionally, the MD-impaired model suffered from cognitive inflexibility, and MD-stimulation could rescue the cognitive deficit (**Fig. 8j**, middle and right panels).

## DISCUSSION

The thalamus has various subnetwork motifs and diverse cortical target outputs, the higher-order thalamic nuclei such as the MD is critical for dynamic regulation of cortical activity in attention, executive control and perceptual decision-making^9,36,37^. Several computational models of thalamocortical networks have been developed for the sensory thalamus^38-40^ and higher-order thalamus^22,29,41^. However, it remains unknown how the MD thalamus contributes to decision making with task uncertainty. Theory and modeling can help unravel key computational mechanisms that provide new experimental predictions^42^. Motivated by multiple lines of animal data^5,6,43^, we trained biologically inspired PFC-MD models with interneuron subtype and thalamocortical pathway specificity to perform two versions of context-dependent decision-making tasks, each with independent sensory and mapping uncertainties. We found that the PFC-MD model outperformed the PFC-alone model in the presence of sensory uncertainty, while cell-type and pathway specificity play a vital role in regulating prefrontal computation during evidence accumulation (cueing period) and working memory (delay period). At different task phases, the MD regulates prefrontal resources for information transmission and maintenance to encode the task variables (rule and context) in a coordinated manner. Rule information is encoded by prefrontal attractor-line dynamics and preserved during the delay period. We found cell-type specific tunings in the MD subpopulation with respect to cueing uncertainty and sparsity, and such MD tunings can be influenced by corticothalamic projection. Additionally, adaptation of MD-PFC connectivity enabled the PFC-MD network to rapidly learn context switching and maintain invariant rule/context encoding, suggesting the role of MD as an ON/OFF switch actuator to enable cognitive flexibility. The notion of thalamus as an ON/OFF switch to assist the thalamocortical control is also in line with the concept in the motor system^44,45^.

Cognitive flexibility requires PFC-MD coordination to map context to behavior, as well as to regulate sensory and mapping uncertainties. Our computational model provides direct support for circuit mechanisms of PFC-MD network in flexible decision making. Distinct cellular targeting among cortical interneurons underlies differential inhibitory effects on excitatory neurons^46^. Cortical interneuron-specific cells that specialize in synaptic disinhibition of excitatory neurons may shape the way excitatory neurons integrate information, and exhibit context-specific and behavior-relevant responses^32,47^. Optogenetic tagging experiments in mice revealed that MD_GRIK4_⟶PV projection suppresses prefrontal noise when task inputs are dense but conflicting, and MD_DRD2_⟶VIP projection amplifies prefrontal signals when task inputs are sparse^5^. The MD_1_⟶PV projection is the primary drive for feedforward thalamic inhibition^48,49^. Cortical PV interneurons have been implied in the mechanisms of gamma oscillations and cognitive function^50^; disturbance in this signaling contributes to altered gamma oscillations and working memory deficits in schizophrenia^51,52^. Thalamocortical and corticothalamic pathways play different roles in decision-making processes^17^. Corticothalamic projection is known to adjust the gain and tuning precision of thalamic neurons as required by behavioral demands^10^. Our analyses of corticothalamic and thalamocortical projections suggest an asymmetric information flow and distinct contributions to decision making in a task phase-dependent manner. The thalamocortical plasticity can potentially modify PFC neuronal correlation^53^, and further improve prefrontal SNR (Methods).

At the computational level, low-dimensional representations of the MD may facilitate compression of cortical information and enable predictive coding in the context of cognitive flexibility. Specifically, since the MD modulates with cue uncertainty and context (“prior”), whereas the PFC encodes the rule (“likelihood”), integrating information from both MD and PFC provides a natural paradigm for updating the posterior in a Bayesian framework^54^.

The capacity of information transmission between neurons or circuits is fundamentally limited by the SNR. Neuromodulation such as dopamine may enhance the SNR of prefrontal activity^55,56^. Since the MD is known to receive dopaminergic inputs^57^, the MD_2_ subpopulation in our model may be viewed as performing a similar function relevant to the dopamine type-2 receptor (D2)-expressing projection^5^. This action of dopamine is achieved by D1 and D2-receptor-mediated effects on pyramidal and local circuit neurons, which further mediate neuronal excitability and recurrent inhibition and thus contribute to the stability of cortical representations of external and internal stimuli^58^. Our results are also consistent with the idea that the MD can facilitate GABA shift and change prefrontal functional connectivity and E/I balance^59^.

Adaptation to rule or behavior according to the task context is a fundamental property in cognitive control. Cognitive flexibility is facilitated by the inclusion of MD^6^—a feedforward low-dimensional structure that monitors both cue conflicts and task error^5^. We reason that adaptive behaviors are driven by both fast and slow learning, where the feedforward structure is appealing to fast learning and the recurrent cortical structure is involved in slow learning. Our theoretical reasoning and computational simulations have shown that thalamocortical plasticity of a pretrained PFC-MD model through trial-by-trial learning enables a rapid and reversible cue-to-rule remapping for a context switch. Moreover, we found that the feedforward thalamocortical connection plays a more vital role than the feedback corticothalamic connections in the speed of context switching. During remapping, rule information is invariantly preserved by PFC population dynamics, resulting in a geometric rotation of neural trajectories.

What computational insight do our models provide to the studies of cognition in a broader context? First, our PFC-MD model provides a means to identify the role of thalamocortical and corticothalamic signaling and how populations of neurons interact between the source and target at different decision-making task phases^60^. Second, computer simulations for various task conditions, which are usually difficult to accomplish in animal experiments due to resource constraints, may produce experimentally testable predictions on behavior and neural representations (such as cortical connection motifs, changes in neuronal tunings under uncertainty). Lastly, our model with cell-type specificity provides a strategy to dissect computational mechanisms of cognitive deficits (e.g., sensory gating and top-down control) encountered in neuropsychiatric disorders such as schizophrenia^61-63^. Abnormal dopaminergic activation and lower cortical SNR have also been implicated in schizophrenia^56^. A lack of inhibitory control increases hyperexcitability and reduces cognitive flexibility in schizophrenia or ADHD^12,64,65^. Dysfunction of GABAergic inhibition may impact synaptic E/I balance that is linked to pathophysiology of a wide range of disease phenotypes such as cognitive deficits and negative symptoms. Recent results have shown that bidirectional plasticity in PFC-MD pathways may correct cognitive impairment^66^. Our modeling results have showed that reduced MD population size or impaired MD⟶PFC pathway reduce the speed of learning the context switch, resembling the cognitive deficits in schizophrenia. Finally, a complete dissection of these computational mechanisms may provide therapeutic insight into thalamic deep brain stimulation strategies to improve cognitive functions in mental disorders.

## Methods

### Context-dependent cognition with parameterized sensory uncertainty

The computational task was modified from a combination of a version of the two-context two-modality four alternative forced choice (4AFC) task^17^ and a single-modality cue-conflicting two alternative force choice (2AFC) task^5^. **Fig. 1a** illustrates the two cueing contexts that were mapped to the same rule: attend-to-audition (rule 1) or attend-to-vision (rule 2). Each trial started with a 40-ms trial initialization period, followed by 800-ms stimulus cueing. In the cueing period, the neural network received cues from either visual or auditory modality, each of which had parameterized cue uncertainty or conflicting cues. In the visual modality context, the random dot motion (RDM) subtask used cues as random dots in motion, where the coherence of moving dots in time indicates the degree of uncertainty: with 1 meaning all dots moving with the same direction (rightward, rule 1 vs. leftward, rule 2) and 0 meaning completely random direction. The default dimensionality of cue input in RDM was 64. In the auditory modality context, the conflicting acoustic cue (CAC) subtask used cues as a sequence of conflicting pulses: low-pass (LP) pulse (+1) or high-pass (HP) pulse (-1). The {+1,-1} pulses were task-relevant cues, where the background was represented by 0. The number of pulses during the 800-ms cueing period was 32 in the training dataset, giving a signal density 40 s^-1^. The mixture of sensory cues determined the uncertainty. Accumulating the sensory evidence (i.e., the total or relative number of LP and HP pulses) would produce the indication of rule 1 (more -1’s) or rule 2 (more +1’s). During the cueing period, the neural network first learned to accumulate the sensory evidence of ambiguous cues to recognize the rule. During the follow-up 800-ms delay period, the neural network learned to preserve the rule information during working memory. During the 200-ms target period, the auditory and visual stimuli (which were used differently from the cueing period) were simultaneously presented (e.g., upsweep/downsweep tone that was mapped to auditory left/right choice, respectively; yellow/blue light that was mapped to visual left/right choice, respectively)^16^, and each attended target was associated with a unique choice. Therefore, the network learned to attend to the correct modality and select one of four choices (**Fig. 1a**). Together, the neural network learned a cross-modal context-dependent working memory and decision-making task, which mapped the ambiguous cueing signals and rules to decision. As a result, the task error could be ascribed to either rule error or sensory error, or both. Specifically, errors related to rule encoding was termed the “rule error” (or “executive error”, which occurred in cueing and delay periods), whereas errors related to target stimulus perception was termed the “sensory error” (which occurred in the post-delay period) (Table S1).

**Table S1.** Categorization of task errors in the context-dependent 4AFC task.

|  | Rule and Context-Dependent Cueing |  |
| --- | --- | --- |
|  | Rule 1: attend-to-audition | Rule 2: attend-to-vision |

| Response \ Target | Cue: A<sub>LP</sub> or V<sub>right</sub> |  | Cue: A<sub>HP</sub> or V<sub>left</sub> |  |
| --- | --- | --- | --- | --- |
|  | Auditory left | Auditory right | Visual left | Visual right |
| Choice #1 (auditory left) | correct | sensory | rule | both |
| Choice #2 (auditory right) | sensory | correct | both | rule |
| Choice #3 (visual left) | rule | both | correct | sensory |
| Choice #4 (visual right) | both | rule | sensory | correct |

In theory, the chance-level accuracy for the 4AFC task is 25%. However, the rule error and sensory error are independent in the task design. For the rule error alone, the chance-level accuracy is 50%. To introduce a proper degree of sensory error in the target period, we assumed two competing sensory inputs were stochastic and drawn from normal distributions with two different mean values but the same variance. Increasing the variance would introduce the task difficulty and increase the sensory error; we selected a variance value that best matched to the experimental data. In the extreme case where sensory inputs were deterministic, the model prediction would lead to only rule error and minimal sensory error.

### Computational models

We first constructed a PFC-alone network as the baseline model. The PFC network is an excitatory-inhibitory (E/I) recurrent neural network (RNN) with *N*_PFC_*=*256 fully interconnected units described by a standard firing-rate model^24,25^. We adopted a continuous-time formulation of RNN dynamics as follows:

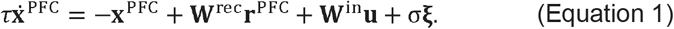

where τ denotes the time constant (we used =20 ms), ξ denotes additive *N*_PFC_-dimensional Gaussian noise, each independently drawn from a standard normal distribution, σ defines the scale of the noise standard deviation; **w**^rec^ is an *N*_PFC_×*N*_PFC_ matrix of recurrent connection weights, and **w**^in^ denotes an *N*_PFC_ ×*N*_in_ matrix of connection weights from the input to network units (*N*_in_ =72). The network output consisted a set of linear readout units that produced a 4-dimensional target estimate: **z** = **w**^out^**r**, where w^out^ is a 4 ×*N*_PFC_ matrix, and the *N*_PFC_-dimensional neuronal firing rate vector is defined by a softplus function: **r** = ϕ(**x**). The network selected the maximum mean output from **z**(*t*) to yield one out of four choices in decision. We assumed 4:1 ratio of the number of excitatory to inhibitory neurons and imposed Dale’s principle on synaptic connectivity. Therefore, the recurrent weight matrix consisted of functionally distinct connectivity according to the cell types: excitatory-to-excitatory, inhibitory-to-excitatory, excitatory-to-inhibitory, and inhibitory-to-inhibitory connections. Among cortical inhibitory neurons, we further considered three major neuron subtypes: parvalbumin (PV), vasoactive intestinal peptide (VIP)-expressing and somatostatin (SOM) interneurons. The interneuron cell types had distinct inhibitory-to-excitatory and inhibitory-to-inhibitory connectivity.^26-29^ The most common PV interneurons target excitatory neuronal cell bodies, and the second most common SOM interneurons target the distal dendrites of postsynaptic excitatory neurons. VIP interneurons preferentially target SOM cells, and to a lesser extent PV cells, so we did not consider VIP⟶PV connectivity here; therefore, VIP interneurons can modulate cortical excitatory neurons through disinhibition of SOM cells (**Fig. 1b**). We further assumed that the percentages of three interneuron cell types were equal and these subtypes were determined by the synaptic interconnectivity. To discretize continuous-time dynamics, we used a bin size of Δ1=20 ms in all numerical simulations.

We further constructed a PFC-MD model based on the knowledge of genetically identified thalamocortical projections and cell type specificity. In light of experimental findings^4^, we assumed that the MD primarily consists of non-recurrent excitatory neurons (*N*_MD_=200) and have two specific MD⟶PFC pathways: where the MD_1_ subpopulation projects to cortical PV interneurons, whereas the MD_2_ subpopulation projects to cortical VIP interneurons. Meanwhile, MD neurons received reciprocal projections from PFC excitatory neurons. We assumed an equal number of excitatory units in both MD_1_ and MD_2_ subpopulations. The MD and PFC units both received a *N*_in_-dimensional stimulus input, but only the PFC units were used to generate a 4-dimensional choice output. There were fully connected corticothalamic connections (of size *N*_PFC_×*N*_MD_) from PFC units to MD_1_ and MD_2_ subpopulation units, separately. For convenience, we let **w**^eff^ represent the effective PFC-MD connectivity matrix of size (*N*_PFC_+*N*_MD_)×(*N*_PFC_+*N*_MD_). In this case, the prefrontal dynamics were controlled by both intracortical weights and thalamocortical weights^44^

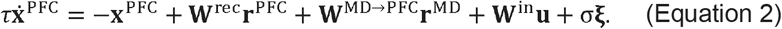

And the MD dynamics were controlled by corticothalamic weights:

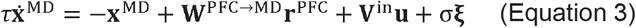

where **v**^in^ denotes an *N*_MD_ ×*N*_in_ matrix. Together, we can rewrite Equations 2 and 3 with a unified equation:

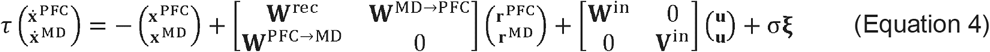

The output setup was identical to the PFC-alone model, where the readout only depended on the PFC units but not MD units.

We trained the PFC-alone and PFC-MD models by supervised learning. The two subtasks, CAC and RDM tasks, were learned separately or simultaneously. The CAC or RDM subtask consisted of a batch size of 64 simulated trials used for training. The number of training trials for two rules (attend-to-audition vs. attend-to-vision) was balanced. We used the mean-squared-error (MSE) plus weight decay as the regularized cost function and optimized the model parameters using a gradient-descent algorithm, with default configuration of hyperparameters (learning rate parameter 0.0005; weight decay regularization parameter 0.1). The optimization continued until the desired accuracy was reached. In total, we trained more than 20 independent PFC-MD networks (and an equally matched number of PFC-alone networks), and selected their respective best models to present the results (however, the psychometric curves and neural representations were quite robust among most task-optimized models). In testing, we varied the degree of cue uncertainty and simulated the 10 independent random test trials per condition in order to compute the psychometric curve. In the CAC subtask, we changed the ratio of LP/HP pulses for different cue uncertainties, and varied the timing and order of the LP and HP pulses for a fixed uncertainty (**Fig. 1g**). In the RDM subtask, we varied the percentage of dots moving rightward or leftward for different cue uncertainties, and selected different subsets of dots in time (**Fig. 1j**). We also observed that some task-optimized PFC-MD and PFC-alone models did not show good generalization across all tested cue uncertainty and sparsity conditions (e.g., making more one type of rule-specific error), which we have excluded in the analyses.

### Decision making with mapping uncertainty: switching cue-to-rule transformation in the CAC task

To introduce mapping uncertainty, we considered a switching context task (*Context 1*⟶*Context 2*⟶ *Context 1’*) using the simulated LP/HP cues in the CAC task (with 800-ms cueing period and 600-ms delay period). In *Context 1*, more LP (LP>HP) pulses would encode rule 1: attend-to-audition. In *Context 2*, the rule was reverse: more LP pulses would encode rule 2: attend-to-vision (**Fig. 7a**). The cue and sensory inputs were identical to the previous setup. At the first stage of learning *Context 1*, we trained the PFC-MD model by batch learning. At the second stage for learning the context switching, we froze intracortical connection weights **w**^rec^, and adapted the other model parameters {**w**^MD⟶tP C^,**w**^P C⟶tMD^, **w**^in^, **v**^in^}. on a trial-by-trial basis. After each trial, the correct or error feedback was used and back-propagated to sequentially update the model parameters. We further learned *Context 2* under the new cue-to-rule transformation, and then relearned the *Context 1’* based on the original rule. To speed up computation, we used a smaller size of network, with *N*_PFC_*=*256 and *N*_MD_*=*30 (*N*_MD1_=15, *N*_MD2_=15). Additionally, we used a gradually annealing learning rule during online learning.

### Behavioral analysis and psychometric curve

The model performance was assessed by a psychometric curve, where the accuracy (percentage of correct) was calculated with varying degrees of cue uncertainty. In the CAC task, the LP/HP congruence level (range: [0.5,1.0]) was defined by a ratio: 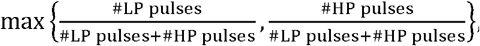, with 0.5 being maximally uncertain and 1 being maximally certain. In the RDM task, the cue uncertainty was calculated by the degree of coherence (range: [-1.0, +1.0]) of random dot motion (with absolute coherence of 1 indicating maximal certainty and noise-free, and coherence of 0 being completely uncertain). Furthermore, we used the algebraic sign to represent the motion direction (with positive/negative values moving rightward/leftward, respectively).^24^ In testing, we generated 10 random realizations of input sequences and calculated the task errors for producing performance curves (mean ± s.e.m.). Based on the percentage of correct choices, we further fit a sigmoid-shaped psychometric curve function using maximum likelihood estimation (**Fig. 1e**). Note that in all psychometric curves, only the rightmost cue condition in the x-axis (i.e., lowest cue uncertainty) was used for training the models, and the remaining cue conditions were never used in training.

### Single-unit tuning curve analysis

We computed the trial-averaged peri-stimulus time histogram (PSTH) from single unit firing activity for specific task condition (rule/context/cue uncertainty/cue sparsity). We excluded the units with very low firing rates (lower 5-10 percentiles of population) in tuning curve analysis. We identified rule or context modulation by comparing their respective mean firing rate as well as temporal PSTH profiles during both cueing and delay periods (**Fig. 2a-e**). Across trials, mean±SD firing rates of units were calculated, the rank-sum test was used to compare the statistics to determine significant modulation (p<0.05). To define the cue uncertainty tuning, we varied the cue coherence or congruence level (high/medium/low) (**Fig. 2f-h**) and compared the respective PSTH profiles. A unit was called to have cue uncertainty modulation if and only if the PSTH comparisons were significantly different between all three paired cue uncertainty levels (high vs. medium, medium vs. low, high vs. low) during the cueing period.

### Excitatory/inhibition (E/I) input

For each PFC excitatory unit, we compute the total excitatory input and inhibitory input

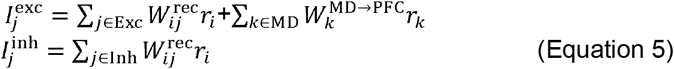

where the total excitatory input consisted of excitatory units of both cortical and thalamic contributions, and the total inhibitory input consisted of input from cortical inhibitory units.

### Neural sequence analysis

To examine potential neural sequences during the delay period, we normalized the PSTHs of PFC excitatory units between 0 and 1 and sorted by the latency of their peak firing rates. Units with peak firing rate smaller than 0.2 during the delay period were excluded in this analysis. We compared the neural sequence among the selected subpopulation between rules and between contexts using the same rank order (**Fig. 2i**).

### Population decoding analysis

We employed a binary support vector machine (SVM) classifier to achieve trial-by-trial classification to decode the task rule or context, based on the simulated population activity of PFC or MD units. The software was implemented under the Sciket-learn Python environment. The network received an *N*-by-1 input, with each representing the mean firing activity of one unit. Among the total 200 simulated trials, we divided them into halves: 50% for training and remaining 50% for testing. We reported the average cross-validation decoding accuracy based on 20 independent Monte Carlo runs. We conducted with decoding analysis under various cue uncertainty conditions (**Fig. 2j, k**).

### Dimensionality reduction analysis

To extract low-dimensional representation of population activity from the PFC-MD network, we employed principal component analysis (PCA) to visualize neural trajectories during the cueing and delay periods. Briefly, we first defined the task-related axes and grouped neural population activity into a matrix **x** 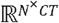, where *N* denotes the number of units, *C* denotes the number of stimulus conditions, and *T* denotes the number of time bins. We performed PCA separately on temporally-binned data matrix **x** at specific or combined task periods, and further extracted two dominant principal subspaces associated with the two largest eigenvalues. The derived low-dimensional neural trajectory showed task-variable-nonspecific representations.

To further visualize the neural trajectory in task-variable-axes (e.g., choice of rule, cue, uncertainty), we further defined the task-relevant state space using a published method^27^. First, we used the 20 largest principal components (PCs) and denoised the representation of neural activity. Second, we constructed a regression model for each unit based on task variables for two context-dependent subtasks: cue and choice of rule. The regression coefficients across units defined an axis in the neural state space representing each task variable. After performing QR orthogonalization, these axes formed the task-relevant state space onto which the population response in the 20-dimensional PC subspace was projected, providing a denoised and demixed task-relevant neural representation. Unless stated otherwise, we used only correct trials to produce trial-averaged curves in two-dimensional subspace (**Fig. 3a-f**). From a neural trajectory ***P*** (*t*), we further defined the time-varying neural velocity by computing the change rate in neural trajectory: 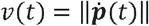 (**Fig. 3g, h**).

### Modeling phasic changes in MD neuronal firing

To model optogenetic activation or suppression effect on the MD neuronal firing, we introduced a depolarizing or hyperpolarizing single-pulse input to the targeted MD neurons, which led to an increase or decrease in their phasic firing activities, respectively (**Fig**. 6b-d).

### Fixed-point analysis of recurrent network dynamics

Similar to our previous analysis^24,25^, we identified fixed-points or slow points of the task-optimized PFC-MD model by numerically solving the optimization problem (https://github.com/mattgolub/fixed-point-finder)

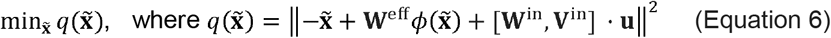

where.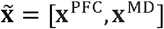 We collected a set of fixed-points by randomly initializing the network. The dimensionality of fixed points was the same as dim 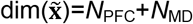. Once the numerical optimization was completed, we applied PCA to visualize the fixed points in three-dimensional PC subspace (**Fig. 3i**).

### Eigenvalue analysis and time constant estimation

A square matrix A is non-normal if it satisfies **AA**^T^ ≠ **A**^T^**A** . For the non-normal matrix 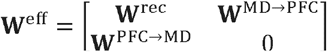 (in the PFC-MD model) or **w**^rec^ (in the MD-lesioned model), we conducted eigenvalue analysis to analyze the nonlinear dynamics with linear approximation^25,67^, where the maximum of the real (or real-part) eigenvalues characterizes the long-term behavior for the speed of network’s steady state decaying to zero^68^. We further we adapted a published method to estimate the intrinsic time constant of dynamical systems^69^, for both PFC-MD and MD-lesioned structures (**Fig. 8d**):

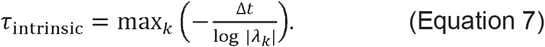

where |λ_k_| denotes the absolute value of the *k*-th real or complex-valued eigenvalue. Since a large value of time constant is beneficial for temporal summation, a larger time constant can help information integration (during the cueing period) and working memory maintenance (during the delay period).

### Mathematical analysis of cue-to-rule transformation

In light of Equation 2 and Equation 4, we assumed that the PFC-MD network converged to a steady state (i.e., the velocity of the state, 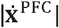 and 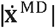 approached to zeros towards the end of task delay period, see **Fig. 3g, h**). To gain some mathematical intuition, we assumed a linear RNN dynamics and let 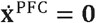 and 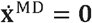. Upon rearranging the terms and assumingτ = *dt*, we derived the steady-state solutions 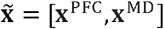:

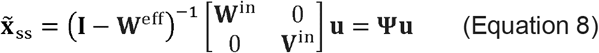

where the (*N*_PFC_+*N*_MD_)-dimensional matrix 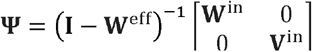 denotes the cue-to-rule transformation. Let **Ψ**_context 1_ and **Ψ**_context 1_ denote the transformation matrices under *Context 1* and *Context 1’*, respectively; if the cue-to-rule tunings of PFC units are reversibly remapped, we will expect that these two transformation matrices are similar (**Supplementary Fig. 6d**). Note that if **u** is univariate, **Ψ** will reduce to a (*N*_PFC_+*N*_MD_)-dimensional vector.

From Equation 2, we see that feedforward MD input and cue input jointly influence the prefrontal neural dynamics 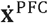 In the two-dimensional PCA subspace, the MD input can shift the original prefrontal dynamics and modify the vector field (**Fig. 8f**). Additionally, the cue input can modulate the MD activity through **v**^in^ (such that the MD can encode cue uncertainty or sparsity). When the MD input is disconnected or disrupted, the cue-to-rule transformation Ψ will be written as **Ψ** = (**I**-**w** ^rec^)^-1^**w**^in^. If the prefrontal internal connectivity is unchanged in the absence of MD-to-PFC feedforward control, the only degree-of-freedom in modulation is through the input weights **w**^in^.

### Optimal thalamic control

The MD thalamus can be viewed as a controller that operates on the cortical plant, which enables the context switch. To gain mathematical insight, we assume that the controlled cortical state in *Context 2* at a fixed point (denoted by **X**^*,PFC, context2^) was a rotated version of state in *Context 1* at the fixed point (denoted by **X**^*,PFC, context1^; see a two-dimensional cartoon illustration in **Fig. 8h**) during the steady state (where 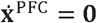 and external input **u**=**0**). From Equation 2, the naïve feedforward strategy to achieve the cortical target state to set the net MD-to-PFC input as^45^

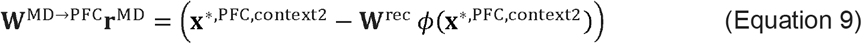

Provided that the intracortical connection **w**^rec^ is intact, then the one-shot feedforward control strategy to learn the context switch (*Context 1* ⟶ *Context 2*) is to set **r**^MD^ as

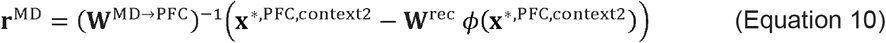

Alternatively, **r**^MD^ can be updated at a slower timescale by modifying the thalamocortical weights **w**^MD ⟶PFC, context1^as well as the corticothalamic weights **w**^MD⟶PFC, context1^ under *Context 1*. This is relatively simple since there is no recurrent weights within the feedforward MD structure. The speed of switching between two contexts depends on the degree-of-freedom of the PFC-MD system and the initial cortical state.

### Intracortical connection motifs and correlation

Cortical networks are known to contain some connectivity motifs.^70^ Let us consider a recurrent network with arbitrary connectivity among *N* PFC neurons. Spike activities of these PFC neurons, denoted by {s(t)} = {[s_1_(t),s_2_(t),…s_N_(t)]}were modeled as inhomogeneous Poisson processes with time-varying rates **Λ** (t) = [λ_1_(t), λ_2_(t), …,λ_N_(t)] and respective baseline rate 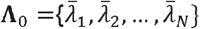 We further modeled the dynamic evolution of **Λ** (t) by a matrix equation:

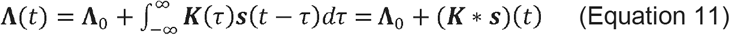

where the effect of presynaptic spikes at time t -τon postsynaptic rates is characterized by the interaction kernels in the matrix **K**(τ) and depends on the elapsed time τ. In the steady state where the expectation value of the rate λ_I_ (t) was not dependent on time (i.e.., E[**Λ** (t)] = **Λ**), we approximated Equation 11 as 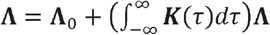 or 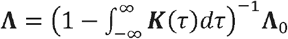. Let 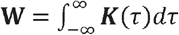, In light of the linear response theory, we rewrote the cross-correlation of neuronal population activity as **C**= (**I**-**W**) ^-1^diag (**Λ**_0_) (**I**-**W**)^-T^. Based on the matrix inverse property (**I**-**W**)^-1^= 1+**w** +**w**^2^ **+…**, without loss of generality, we assumed the rate _0_ as an all-one vector and derived an approximate form of using the first four lower-order expansion terms:

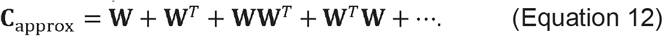

Therefore, the correlation of neuronal activity is influenced by the sum of a set of connection motifs. Replacing **W** of Equation 12 with **w**^eff^, and changing MD presynaptic connectivity **w**^MD → PFC^ provides a way to test how presynaptic feedforward MD input may affect postsynaptic recurrent PFC connection motifs and correlation.

## Data and code availability

All the data that supports the plots within this paper and other findings of this study are available. (https://github.com/Xh-Zhang1/pfc-md).

## Acknowledgements

We thank N. Lam, A. Mukherjee, R.D. Wimmer and Y. Liu for valuable comments on the manuscript. The work was supported by grants MH118928 (Z.S.C.), DA056394 (Z.S.C.), MH132642 (Z.S.C. and M.M.H.) from the US National Institutes of Health. We thank the Conte Center Team for scientific input and inspiration on cognitive thalamus. A preliminary version of this work has appeared in BioRxiv preprint (https://biorxiv.org/cgi/content/short/2022.12.11.519975v1) and was presented in COSYNE’23.

## Author contributions

Z.S.C. conceived and supervised experiments, analyzed and interpreted the data, and wrote the paper. X.Z. developed the computational models, performed experiments, analyzed and interpreted the data. M.M.H. provided experimental data and interpreted the data. Z.S.C. acquired funding.

