## Supplementary material for "Mediodorsal thalamus regulates sensory and mapping uncertainties in flexible decision making": Supp Figures

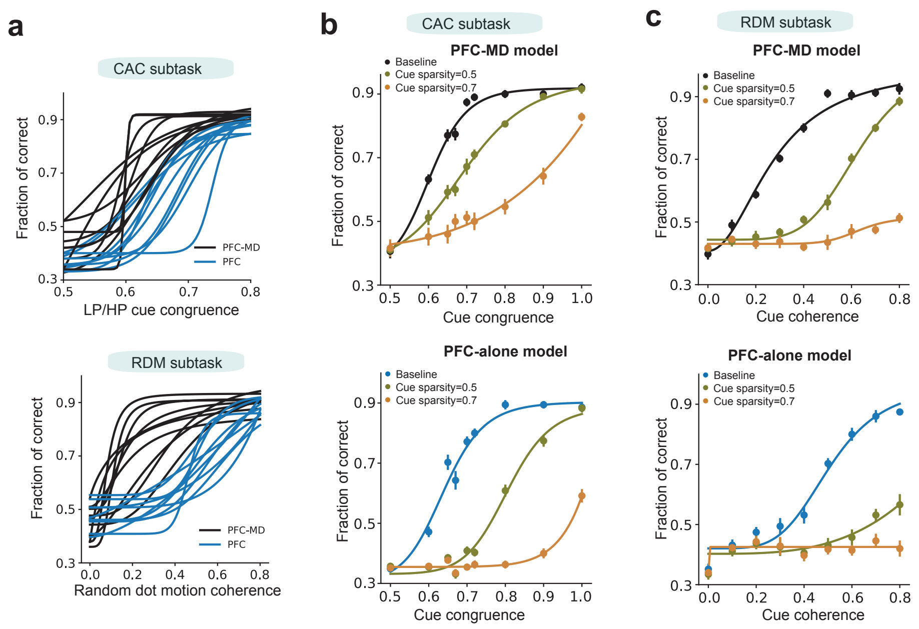


**Supplementary Figure 1**. **Psychometric curve comparison between PFC-MD and PFC-alone models that were both performance-optimized.**
**a,** Selected psychometric curves of PFC-MD and PFC-alone models for both CAC and RDM subtasks. **b,** Psychometric curves of PFC-MD and PFC-alone models for three levels of cue sparsity in the CAC subtask. Cue sparsity 0.0 denotes baseline. **c,** Psychometric curves of PFC-MD and PFC-alone models for three levels of cue sparsity in the RDM subtask.

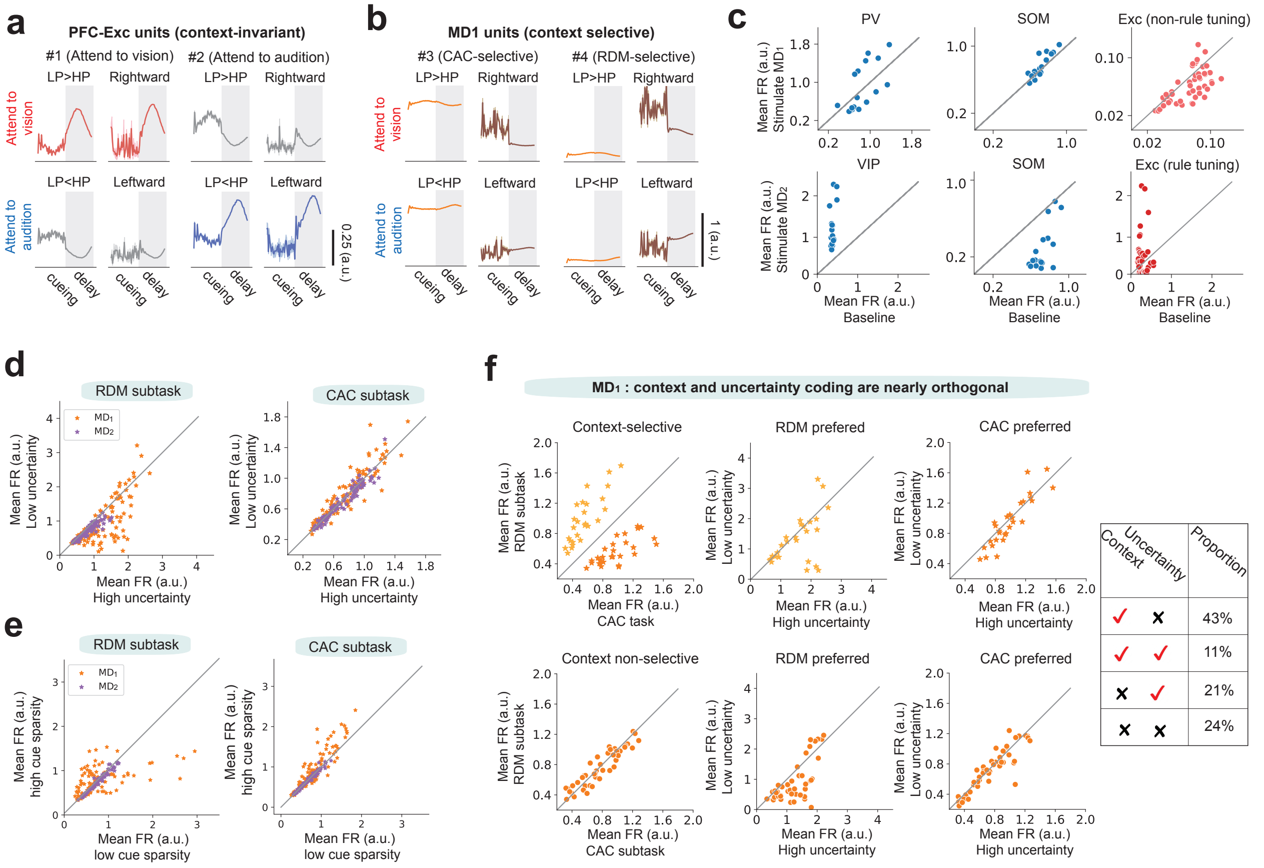


**Supplementary Figure 2**. **PFC and MD single-unit modulation in the performance-optimized PFC-MD model.**
**a,** Two additional examples (similar to **Fig. 2a**) of context-invariant PFC excitatory units encoding two rules under two contexts. **b,** Two additional examples (similar to **Fig. 2b**) of context-selective PFC inhibitory units encoding the cue context. **c,** Two additional examples (similar to **Fig. 2c**) of context-selective MD units encoding context information. **d,** The examples of MD$_2$ units showed no firing modulation with respect to cue sparsity. This was in contrast to MD$_1$ units shown in **Fig. 2h**. **e,** Population statistics of mean firing rates (during the cueing period) of MD$_1$ and MD$_2$ units for encoding cue uncertainty and cue sparsity.

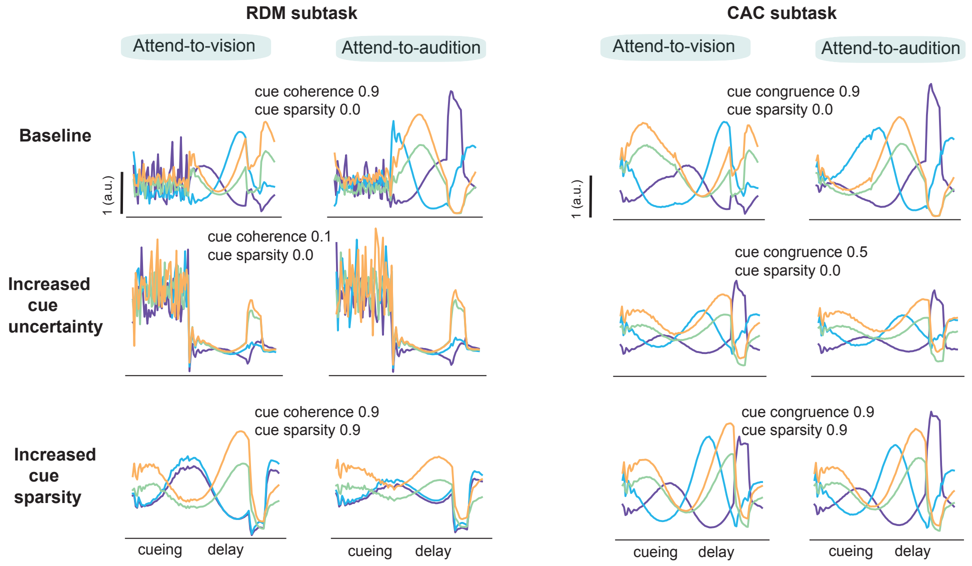


**Supplementary Figure 3. Demonstrations of PFC excitatory units that reduce rule tuning selectivity with increasing cue uncertainty or cue sparsity.**

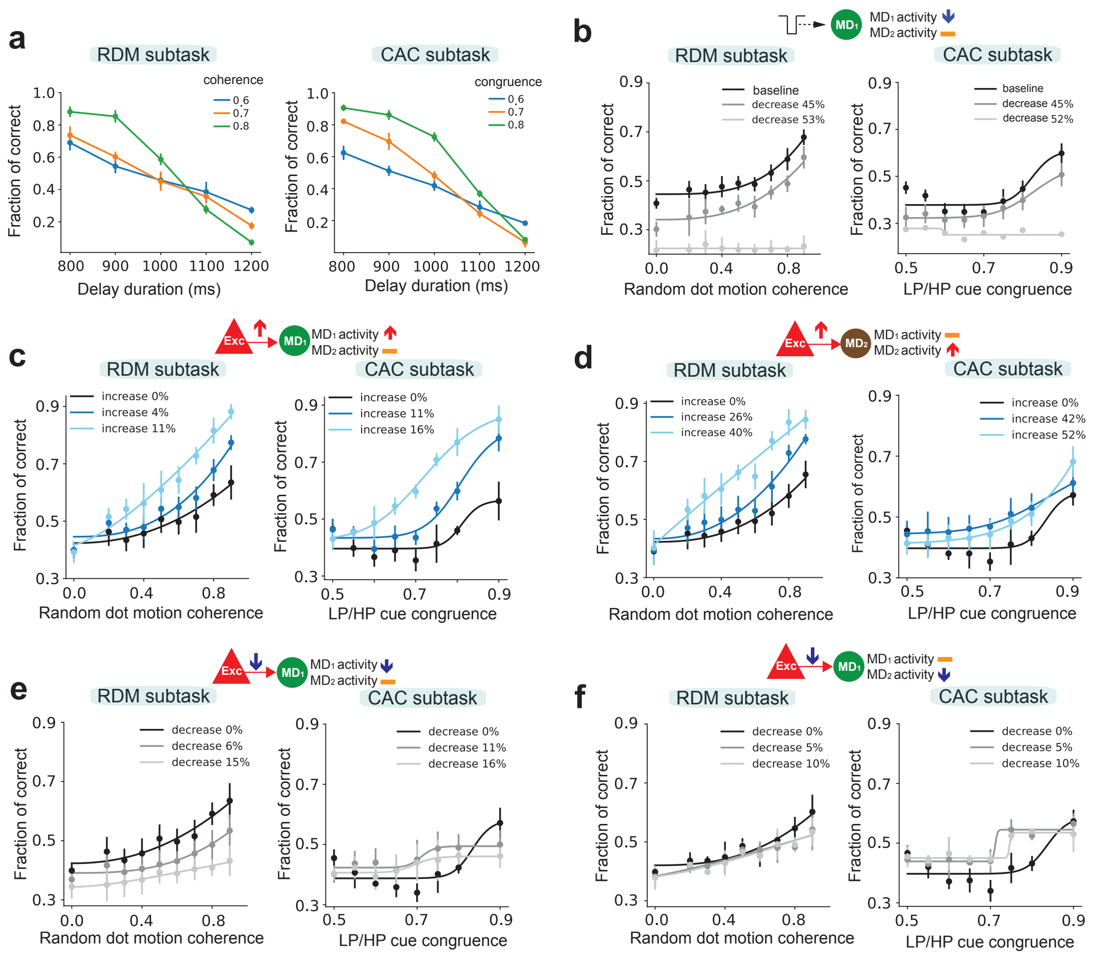


**Supplementary Figure 4. Changing directed prefrontal Exc→MD connection strengths of the PFC-MD model affects MD activity and task performance.**
**a,** The PFC-alone model degraded in accuracy when the delay duration was increased. Error bar denotes SD ($n = 10$). Figure legend denotes the level of cue coherence or cue congruence. **b,** Decreasing the $MD_1$ population firing rate during an elongated delay period had an opposite effect on the task performance. **c,** Strengthening PFC→$MD_1$ connection strengths (shown by a relative increase with respect to the baseline) increased the $MD_1$ firing rate while keeping the $MD_2$ firing rate unchanged, which further boosted working memory maintenance or improved task performance with elongated delay duration. **d,** Strengthening PFC→$MD_2$ connection strengths increased the $MD_2$ activity while keeping the $MD_1$ activity unchanged, which further boosted working memory maintenance during elongated delay. **e,f,** In contrast, weakening PFC→$MD_1$ or PFC→$MD_2$ connection strengths (shown by a relative decrease with respect to the baseline) decreased the respective MD activity, causing a degraded or inconsistent task performance.

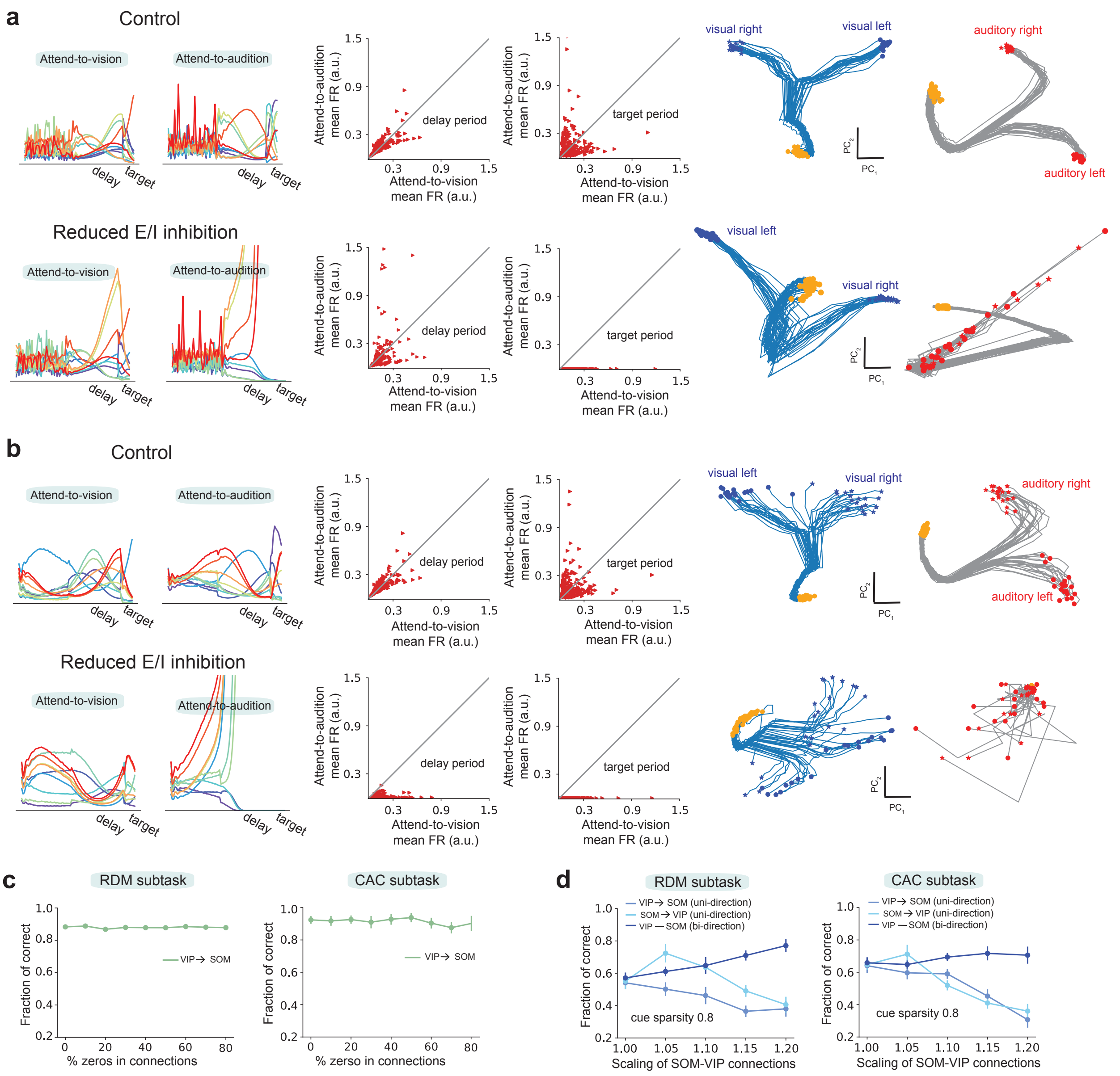


**Supplementary Figure 5**. **Circuit dissection of cognitive deficits.** Comparisons of prefrontal excitatory single-unit tunings, mean firing rates of excitatory population, and low-dimensional neural trajectories between control and reduced E/I inhibition (by changing prefrontal VIP→Exc connectivity) conditions, for RDM subtask **(a)** and CAC subtask **(b)**. **c,**Weakening VIP→SOM connectivity of the performance-optimized PFC-MD model had little impact on task performance. **d,** When the cue signal was sparse, bidirectional SOM-VIP amplification of mutual inhibition strengths could amplify the sparse cue signal, but similar unidirectional manipulations were ineffective.

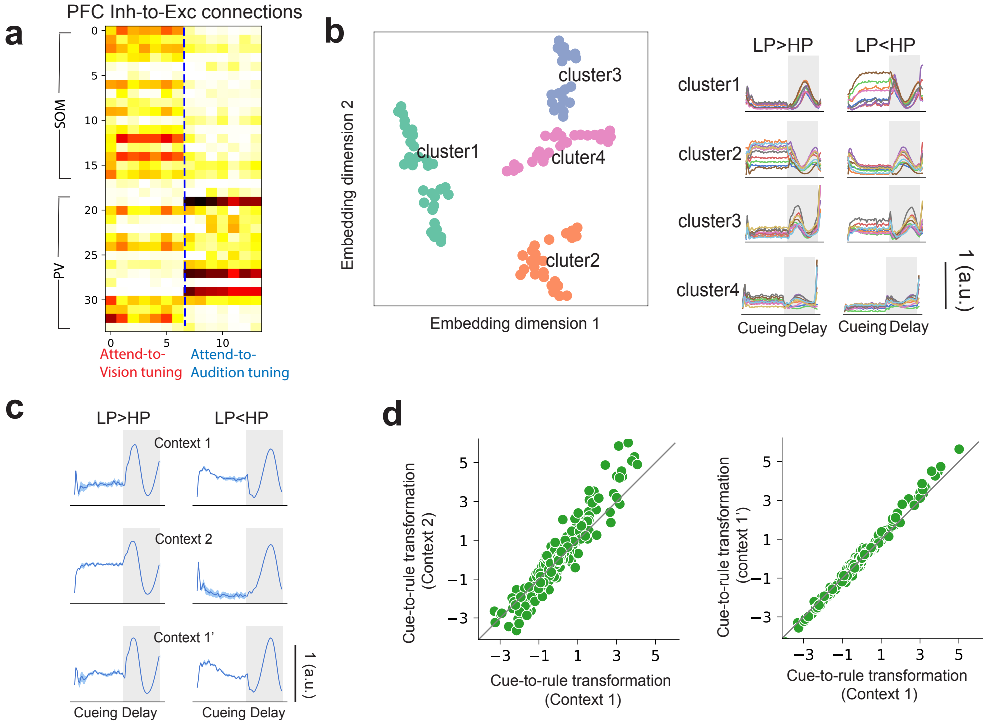


**Supplementary Figure 6**. **The prefrontal activity in context-switching.**
**a,** PFC inhibitory-to-excitatory (Inh-to-Exc) connectivity was mapped into two classes of rule-tuned excitatory units (attend-to-vision vs. attend-to-audition). Only 14 representative PFC units were shown. **b,** PFC excitatory units ($n = 205$) were clustered in two-dimensional space by applying t-SNE embedding to combined Inh-to-Exc, Exc-to-Exc and MD-to-Exc synaptic weights. Four functional cell-type clusters emerged, each with a distinct rule-tuning profile. **c,** Some PFC excitatory units shifted their peak firing rates temporally during the delay period during context switching. **d,** Quantification and comparison of cue-to-rule transformation in prefrontal computation during context switching. Each dot corresponded to the gain related to each PFC unit ($n = 250$) (see Methods).
